# ANP32E drives vulnerability to ATR inhibitors by inducing R-loops-dependent Transcription Replication Conflicts in Triple Negative Breast Cancer

**DOI:** 10.1101/2024.03.25.586539

**Authors:** Sara Lago, Vittoria Poli, Lisa Fol, Mattia Botteon, Alessandra Fasciani, Alice Turdo, Miriam Gaggianesi, Matilde Todaro, Yari Ciani, Giacomo D’Amato, Francesca Demichelis, Alessio Zippo

## Abstract

Oncogene-induced replicative stress (RS) plays a central role in tumor progression, leading to genomic instability by eliciting transcription replication conflicts (TRCs), which represent the major source of R-loops, that ultimately favors the onset of the DNA damage response (DDR). We investigated the pathogenic contribution of chromatin factors in increasing TRCs and R-loop frequencies in cancer. We found that in breast cancer patients the concomitant upregulation of MYC and the H2A.Z-specific chaperone ANP32E correlated with an increase genome instability. Genome-wide profiling revealed that the ANP32E-dependent increases turnover of H2A.Z altered RNApol II processivity, leading to accumulation of long R-loops at TRCs. We showed that ANP32E upregulation increases TRCs and activates an ATR-dependent DDR, which predispose cancer cells to R-loop-mediated genomic fragility. By exploiting the vulnerability of ANP32E-expressing cancer cells to ATR inhibitors, we found that tumors relied on this DDR pathway, whose inhibition halted their pro-metastatic capacity.

## Introduction

Triple negative breast cancer (TNBC) is the most aggressive breast cancer (BC) subtype due to its high recurrence rates, metastatic potential, and heterogeneity.^1^. One determinant of the molecular variability within tumors is genomic instability, eliciting the adaptation and survival capability of pro-metastatic cancer cells, once challenged by hostile microenvironments. Genomic instability can manifest in different forms, like increased frequency of point mutations, copy number alterations (CNA) or chromosomal instability (CIN), which are characterized by changes in the chromosome structure and number over time^2^. Although the underlying molecular processes are still a field of exploration, it has been established that oncogene-induced replicative stress (RS)^3^ plays a central role in tumor progression. For instance, MYC is one of the oncogenes that drives RS in many tumors, and is the most frequently amplified gene in TNBC^4–6^. Some genomic regions, known as Common Fragile Sites (CFSs), are susceptible to RS and to acquiring genomic instability^7^. Despite eukaryotic cells having evolved several mechanisms to keep spatial and temporal separation between transcription and replication, RS may inevitably lead to transcription-replication conflicts (TRC). TRC, hence the collision between the two machineries, is often associated with DNA damage and the formation of genomic lesions^8,9^. This implies that if the TRC is not resolved and/or the DNA damage response (DDR) is not properly executed, they lead to DNA double strand breaks (DSBs) and ultimately to genome instability^10^. Accordingly, cancer cells harboring frequent TRC rely on DNA repair pathways to withstand these genomic challenges, thus representing a synthetic lethality and providing an opportunity for therapeutic intervention^11^. Cancer patients with DDR-addicted tumors would therefore benefit from selective DDR inhibition.

DNA damage resulting from TRC is further worsened by R-loops, which are co-transcriptional DNA:RNA hybrids^12^. In particular, R-loops may favor TRC by forming stable complexes with RNApol II, leading to transcriptional block along the gene body-RNApol II stalling-, or by impeding replicative forks restart at sites of fork stalling^12,13^. Recently, a complex interplay between R-loops and many chromatin factors has been shown to contribute to the control of RNApol II density and processivity^14,15^. Indeed, R-loops play a role in the tight control of RNApol II promoter-proximal pause-release^16^, yet the first barrier in determining the RNApol II promoter escape and processivity along the gene body is represented by nucleosome composition and positioning^17^. The latters are defined by the action of specific chromatin factors, among which histone chaperones, that participate in different steps of nucleosome assembly and recycling^18^. For instance, H2A.X and H2A.Z variants of histone H2A are exchanged by specific chaperones at sites of DNA damage and in the proximity of the transcription start site (TSS), respectively^19^. In particular, H2A.Z is enriched at the +1 nucleosomes, and its turnover is critical to modulate the RNApol II pause-release ^20–22^. Based on their key regulatory functions, we hypostatized that altering chromatin players could amplify oncogene-induced RS beneath genome instability, by potentially increasing TRC and R-loop frequencies.

In the present work, we found that ANP32E is the most frequent amplified and overexpressed histone chaperone in TNBC patients characterized by MYC deregulation. ANP32E chaperone activity is directed towards the eviction of H2A.Z histone variant^23,24^. This study investigated how ANP32E overexpression can worsen the effects of oncogene-induced RS by stimulating TRC, R-loops, and genomic instability, and the therapeutic implications of this mechanism.

## Results

### ANP32E expression correlates with MYC and genome instability in TNBC

To identify histone chaperones that may drive genomic instability in TNBC undergoing MYC-induced RS, we investigated public datasets from BC patients. Given the positive correlation between MYC expression and its genomic amplification in Basal BC (mostly TNBC) (Extended Data Fig.1a)^25^, we looked for histone-binding proteins whose expression is upregulated in MYC-amplified Basal BC patients, when compared to the other BC subtypes. Among the chromatin binding factors, the histone chaperone ANP32E resulted between the most significantly upregulated genes (Fig. 1a, Supplementary Table 1). Importantly, we observed that ANP32E is among the most frequently amplified genes in Basal BC patients and that its expression correlates with higher accumulation and frequency of Copy Number Alterations (CNA), suggesting a potential clinical relevance (Fig. 1b and Extended Data Fig 1b.). To investigate a possible synergistic effect of MYC and ANP32E deregulation in promoting genome instability, we segregated patients based on MYC and ANP32E overexpression matched by their respective copy number gain status in all BC patients (thereafter indicated as MYC-ANP32E). We observed a significant increase in the fraction of altered genome, indicative of genome instability, in patients with the concomitant upregulation of both MYC and ANP32E (Fig. 1c). We also found that the co-amplification of MYC and ANP32E was more frequent in Basal BC with respect to the other BC subtypes (Extended Data Fig.1c). Basal BCs are the most affected by genome instability, a notion that is confirmed by the higher mutation counts, fraction of altered genome, and microsatellite instability (MSI) in comparison with the other BC subtypes (Extended Data Fig.1d-f)^26^. In order to pursue the correlation between MYC-ANP32E upregulation and genome instability, we employed data from a study (Metabric), which clustered patients based on cancer driver CNA influencing gene expression^27,28^. Interestingly we observed that Cluster 9 and Cluster 10, resembling genomically unstable ER+ and Basal TNBC patients respectively, were characterized by the highest frequency of MYC-ANP32E co-upregulation (Fig. 1d and Extended Data Fig. 1g). This is particularly relevant when comparing Cluster 10 with the other cluster of TNBC patients (Cluster 4ER-), which is instead more genomically stable and characterized by the low frequency of MYC-ANP32E co-upregulation (Fig. 1d and Extended Data Fig. 1g). Although high ANP32E overexpression was already reported as an adverse prognostic factor in TNBC, we also observed that the MYC-ANP32E co-upregulation resulted in a worse prognosis for TNBC patients in different cohorts, even when compared with patients with single MYC or ANP32E upregulation (Fig. 1e and Extended Data Fig. 1h)^29^. Since TNBC mortality is linked to invasiveness and metastatization, we analyzed an additional dataset segregating patients for the presence or absence of metastases (Metastatic Breast Cancer Project 2021). This revealed a higher frequency of MYC-ANP32E co-upregulation in TNBC metastatic patients, suggesting that it might provide a cell growth advantage to pro-metastatic cancer cells (Fig. 1f).

**Figure 1.**
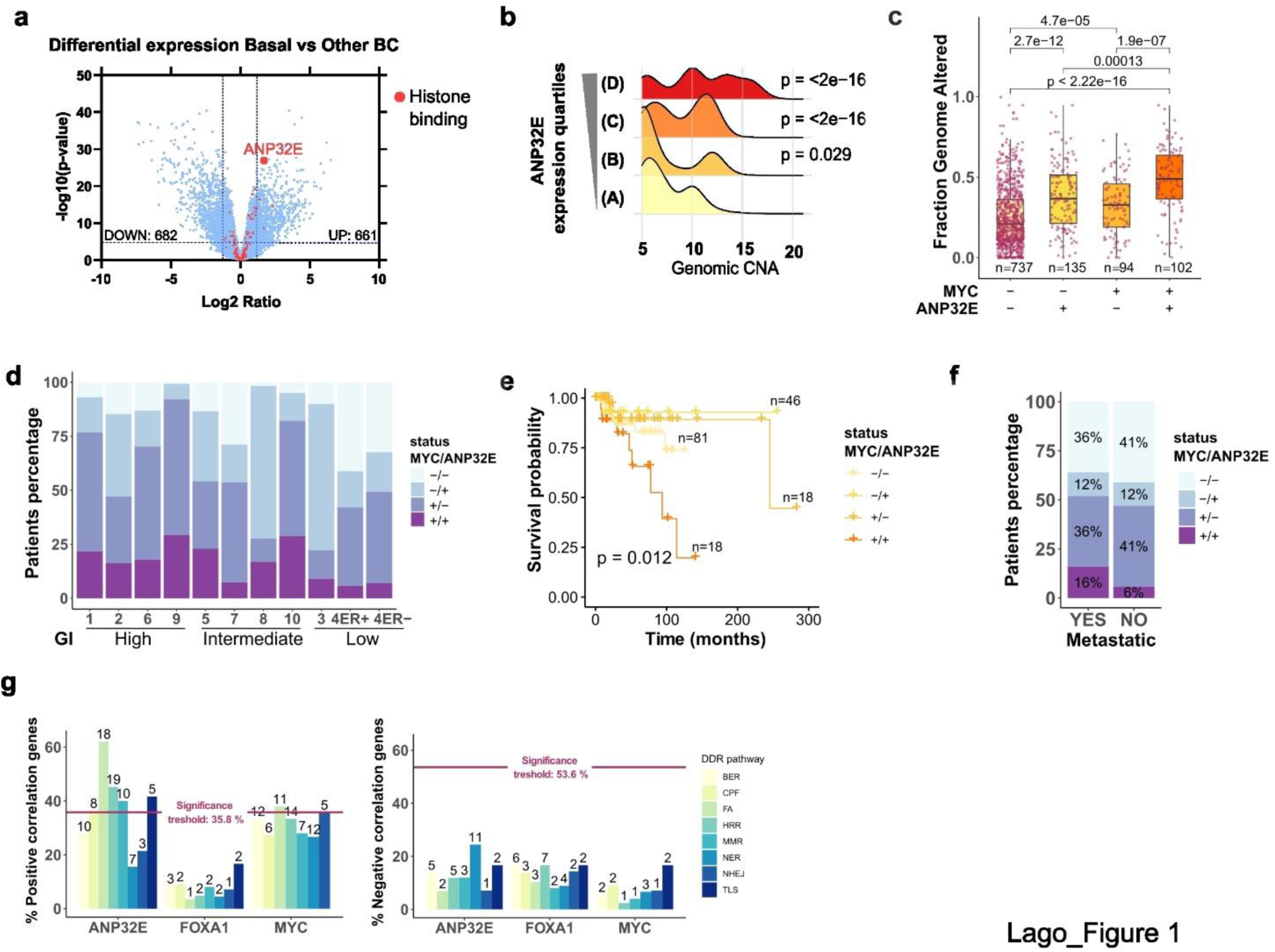
ANP32E expression in Basal BC correlates with genome instability. (**a**) Volcano plot showing differentially expressed genes between MYC-CNA patients with Basal-BC (n=61) or Other-BC subtypes (n=92), retrieved from TCGA data. Chromatin players are shown in red. Dashed lines indicate differential expression significance thresholds: log2FC > ±1.5 and – log10pvalue > 5. Up and downregulated gene numbers are indicated in the graph. **(b)** Density plot showing the frequency of Basal BC patients with genomic CNA, stratified according to ANP32E expression quartiles, expressed as mRNA expression z-scores relative to normal samples (log RNA Seq V2 RSEM). Quartiles: (A) –6.83 – –3.55, (B) –3.56 – 5.13, (C) 5.15 – 6.44, (D) 6.46 – 9.06. Statistical significance measured with t-test. N=171. Data retrieved from TCGA. **(c)** Boxplot showing the fraction of altered genome for all BC patients grouped based on MYC and ANP32E gain and expression. Double positivity was assessed based on 75° of expression for both genes and concomitant copy number gain. Statistical significance measured with t-test. Data retrieved from TCGA. **(d)** Stacked barplot showing the percentage of BC patients with the different combinations of MYC and ANP32E upregulation, stratified according to Metabric IntClust. IntClust are sorted based on the genomic instability (GI) reported level (Low, Intermediate, High). **(e)** Kaplan-Meier plot representing the survival probability of Basal BC patients with the different combinations of ANP32E and MYC upregulation. Data retrieved from TCGA. Double positivity was assessed based on 75° of expression for both genes and concomitant copy number gain. P-values indicate statistical significance for the comparison of groups calculated by log-rank test. **(f)** Barplot reporting the percentage of TNBC patients with the different combinations of MYC and ANP32E co-AMP, grouped according to the presence or absence of metastases. Data retrieved from the Metastatic Breast cancer project 2021. **(g)** Barplot showing the percentage and raw number (displayed above bars) of DDR pathway genes that positively (left panel) or negatively (right panel) correlate with ANP32E expression. Significance threshold calculated by random gene sets analysis is shown. The correlation score was calculated with Pearson method, considering a threshold of +/− 0.85.

Often, genomic instability accumulates in cancers that cannot efficiently repair DNA damage, being deficient in DDR pathways^30–32^. We, therefore, looked at the correlation between ANP32E expression and the transcriptional regulation of specific DDR pathway genes in Basal BC patients, using FOXA1 and MYC expression as negative and positive controls, respectively. We found that several DDR pathway genes show an upregulation correlating with ANP32E expression (Fig. 1g left, Extended Data Fig. 1i). On the contrary, we found no DDR pathway that is downregulated in accordance with ANP32E expression (Fig. 1g right, Extended Data Fig. 1i). The relevance of ANP32E itself in relation to DDR was strengthened by the observation that this association is mostly lost using as reference gene MYC or FOXA1, the latter being inversely associated with hormone receptor-negative BCs^33^. Importantly, we found that the Fanconi Anemia (FA) pathway, involved in resolving RS-derived genome instability, had the strongest correlation with ANP32E deregulation (Fig. 1g left). In sum, this association analysis indicated that in BC patients, the concomitant upregulation of MYC and ANP32E correlated with an increase in genome instability.

### ANP32E elicits R-loop-mediated TRCs and DNA damage

ANP32E-dependent turnover of the histone variant H2A.Z is fundamental for the proper fulfillment of transcription^34^. Deregulated gene transcription can lead to collisions with the DNA replication machinery giving rise to TRCs and DNA damage. Despites its relevance in cancer, a direct link between ANP32E overexpression and TRC was never explored. To this aim, we employed tIMEC cells, a tumorigenic Basal BC cellular model that was previously generated by hTERT-immortalization of human mammary epithelial cells combined with c-MYC and PIK3CA^H1047R^ overexpression^35^. Here we stably overexpressed ANP32E in tIMEC to generate tIMEC-A cells (Extended Data Fig. 2a). As TRCs outcome is the formation of DNA:RNA hybrids, termed R-loops^12,36^, we monitored their accumulation in the designed cellular models. In addition, to test a possible involvement of R-loops, we generated tIMEC-A cells that stably overexpressed RNaseH1 (thereafter named tIMEC-A-H1), which specifically degrades DNA:RNA hybrids (Extended Data Fig. 2b). We found a significant increase of R-loops in tIMEC-A cells, which was rescued by both exogenous treatment with RNaseH1 and its endogenous stable overexpression (Fig. 2a). This result was also confirmed by PLA experiments showing a higher accumulation of R-loops at transcriptionally active sites marked by elongating RNApol II (pS2) (Extended Data Fig. 2c) and in proximity to ANP32E, supporting a direct involvement of ANP32E in R-loops dynamics^37^ (Fig. 2b, d). In addition, TRCs measured as the collision between elongating RNApol II and PCNA, a component of the DNA replication machinery, showed a higher frequency of PLA foci in tIMEC-A cells, which were resolved in tIMEC-A-H1 (Fig. 2c, d). Most probably the conflicts that we visualized are Head-ON conflicts, as only when there is a frontal collision between the transcriptional and replicative machineries the elongating RNApol II and PCNA come in proximity to be detected by PLA^38^.

**Figure 2.**
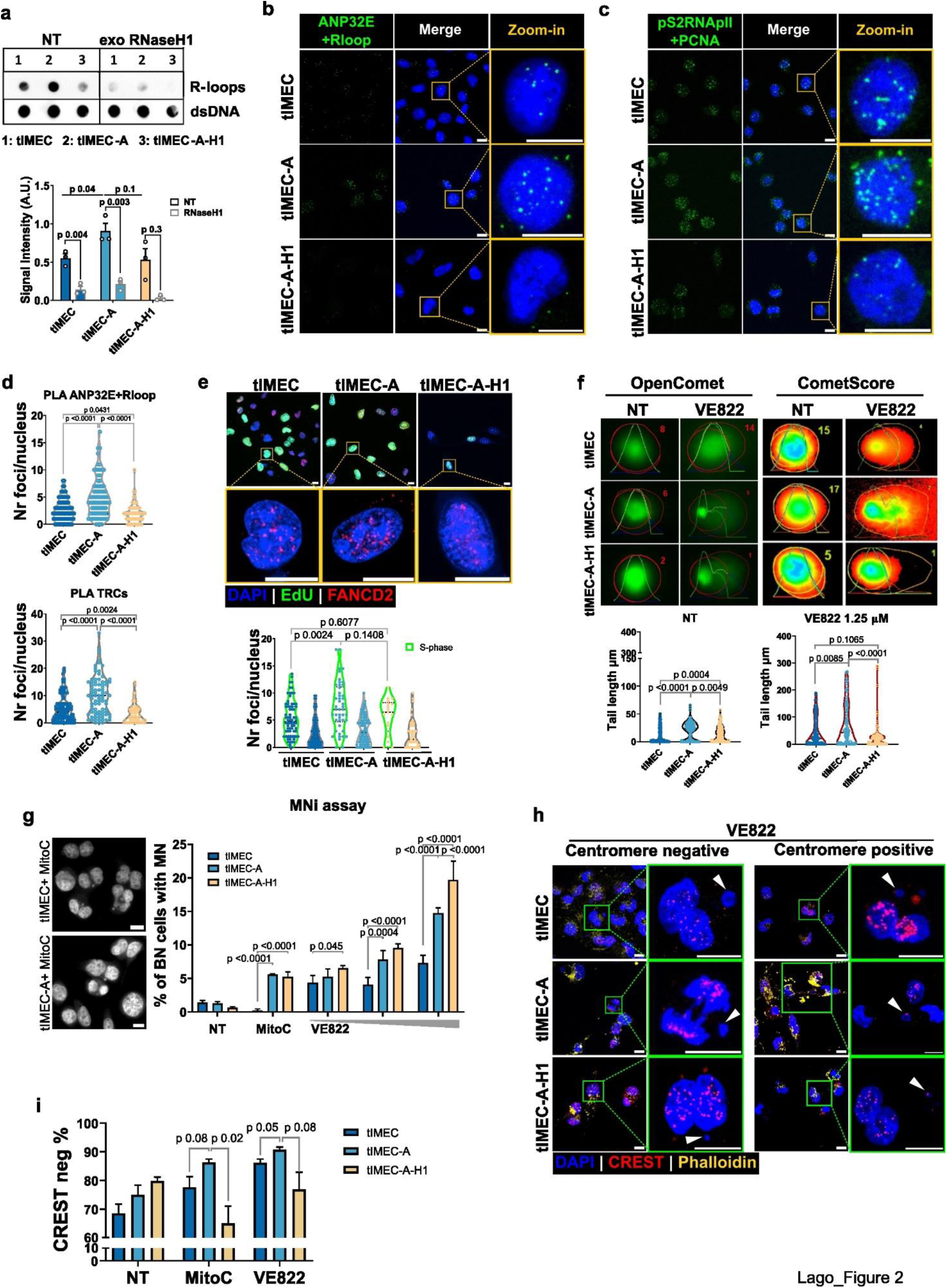
ANP32E overexpression results in increased R-loops, TRCs and DNA damage. (**a**) Top: Dot blot showing the abundance of R-loops in tIMEC, tIMEC-A and tIMEC-A-H1 cell lines with or without exogenous RNaseH1 treatment. dsDNA was measured as internal normalizer. Bottom: Quantification of three replicates with t-test p-value. **(b)** Representative PLA foci images representing proximity between ANP32E and R-loops. Scale bar = 10 µm. **(c)** Representative PLA foci images representing proximity between pS2RNApolII and PCNA as loci for TRCs. Scale bar = 10 µm. **(d)** Quantification of PLA foci per nucleus of experiments reported in (B) (top) and (C) (bottom). The number of measured cells is tIMEC n=106, tIMEC-A n=85, tIMEC-A-H1 n=72, and tIMEC n=104, tIMEC-A n=84, tIMEC-A-H1 n=75, respectively. Unpaired t-test p-values are reported in the graph. Scale bar = 10 µm. **(e)** Top: representative images of FANCD2 immunofluorescence in EdU stained cells. EdU positive cells were chosen as zoom-in but EdU channel was excluded for clarity of representation. Bottom: Quantification of FANCD2 foci per nucleus in EdU + (green violins) or EdU-(gray violins) cells. Unpaired Tttest p-values are reported in the graph. Scale bar = 10 µm. **(f)** Alkaline comet assay performed in the presence or absence of 24h VE822 1.25 µM treatment. Upper panel shows representative comets with quantification metrics for comet head and tail calculated by OpenComet ImageJ plugin^71^ and heatmaps showing the DNA staining intensity measured by CometScore software^72^. The lower panel shows violin plots of comet tail length quantification. 150-300 cells were analyzed for every condition in 3 biological replicates. Unpaired t-test was applied to calculate p-value of statistical significance as reported in graph. **(g)** Left: representative images of binucleated cells (BN) with MNi upon Cytochalasin B + Mitomycin C treatment. Right: percentage of BN cells with MN quantified by Operetta Software dedicated program. Quantifications are reported in the absence or presence of Mitomycin C (0.03 µg/ml) or VE822 (0.027, 0.08, 0.25 µM). The experiment was performed in three biological replicates and at least 3000 cells per condition were measured. The barplot reports mean ± sd with two-way ANOVA adjusted p-values of significant comparisons. **(h)** Representative images of immunostaining on BN cells with MNi that are positive or negative for centromere staining (CREST) in VE822 treated condition. Phalloidin staining was employed to identify cytoplasms. White arrows indicate MNi. Cells were treated as indicated in panel (G) to block cytokinesis. Scale bar = 10 µm. **(i)** Barplot reporting the percentage of BN cells with MNi derived from DSBs (i.e. free of centromeres) in cells prepared according to the procedure described in Supplementary Figure S2E, with or without additional treatment with MitoC or VE822. Mean and s.e.m. of three independent biological replicates are reported. 25-60 BN cells per condition were quantified in every replicate.

Given the observed ANP32E-dependent increase of R-loops and TRCs, we asked whether high ANP32E expression might be a source for DSBs accumulation. Through immunofluorescence analysis, we first noticed that S-phase (EdU+) ANP32E-overexpressing cells showed an increased number of ubiquitinated FANCD2 foci, an ATR-dependent DDR factor that is recruited to the chromatin by binding to R-loops in regions of stalled replication forks (Fig. 2e)^39,40^. Next, we performed an alkaline Comet assay in the presence or absence of two DNA damage stimulating drugs: Etoposide (ETP), a DNA topoisomerases inhibitor that promotes DSBs^41^, and VE822, an ATR inhibitor (ATRi) that impedes ATR-mediated DDR activation^42^. Interestingly, we observed that upon ETP treatment, ANP32E overexpression was sufficient to induce an increase in DSBs in a mechanism likely involving R-loops, since the phenotype was reversed by RNaseH1 upregulation (Extended Data Fig. 2d). Treatment with VE822 led to a sustained increase in DSBs especially in tIMEC-A, suggesting that ANP32E-overexpressing cells strongly rely on DDR activation (Fig. 2f). To test the severity of the accumulated DSBs, we measured the formation of Micronuclei (MNi), small nuclear-like structures containing massively damaged DNA. Despite the absence of a significant difference in untreated cells, both Mitomycin C, used as control clastogenic drug, and VE822 treatments led to a significantly higher increase in MNi formation, especially in ANP32E-overexpressing cells (Fig. 2g and Extended Data Fig. 2e). However, R-loops resolution by RNaseH1 overexpression did not rescue the accumulation of MNi. To gain further insights, we tested whether R-loops are specifically implicated in clastogenic MNi subclass, that derived from unresolved DSBs^43^. Thus, we measured the fraction of MNi containing acentric chromosomes, characterized by the absence of centromeres (CREST-negative). We found that clastogenic MNi are indeed more abundant in cells overexpressing ANP32E and their increase elicited by Mitomycin C or VE822 treatment was both ANP32E and R-loops dependent (Fig. 2h, i, and Extended Data Fig. 2f, g). To exclude that the described mechanism is restricted to the employed cellular model, we selected MDA-MB-231 and SUM-159-PT as two alternative TNBC cell lines that express either high or low levels of ANP32E, respectively, in the presence of upregulated MYC (Extended Data Fig. 3a, b). By either silencing or overexpressing ANP32E in such alternative models (named MB-231^sh^ and SUM-159-A, respectively), we confirmed the correlation between ANP32E modulation, the accumulation of R-loops and TRCs, as well as the tendency to accumulate DNA DSBs (Extended Data Fig. 3c-i). In summary, we found that ANP32E upregulation in TNBC is associated with increased TRCs and DSBs, which predispose cells to genomic fragility via a mechanism involving R-loops. This phenotype is strongly worsened when ATR-mediated DDR is impaired, highlighting a putative vulnerability of BC cells with such biological background.

### ANP32E stimulates ATR-dependent DDR and alteration of transcription and replication dynamics

We observed that the inhibition of ATR in TNBC cells combined with the overexpression of ANP32E generated an augmented genome instability. This result suggested a sustained activation of ATR-mediated DDR in combination with ANP32E expression. We therefore performed immunofluorescence analysis of two key ATR-dependent DDR players, p(S33)RPA32 and 53BP1 (Fig. 3a-d) ^44–48^. We observed a puncta-like distribution of these DDR markers, indicating the activation of the DDR response at discrete genomic loci, that resulted in being more pronounced in tIMEC-A and, depending on ATR signaling as the inhibition of its kinase activity, rescued the ANP32E-dependent increase of DDR. We also observed an increased number of foci-positive cells for both factors in tIMEC-A, indicating that the damage is more spread across the cell population (Fig. 3e). To integrate this information with TRCs we also performed EdU staining to mark cells in S-phase, which is when TRCs are expected to occur. Importantly, we observed that pRPA32 and 53BP1 foci are more abundant in S-phase tIMEC-A cells and rescued with R-loops degradation (Fig. 3a-d). We found that inhibiting ATR by VE822 led to a peculiar phenotype in which nuclei appeared entirely positive to pRPA32 staining, especially in tIMEC-A cells (Fig. 3a, f). The latter condition is attributed to DNA damage catastrophes preceding cell death due to the excessive accumulation of DNA damage^49^. Moreover, despite 53BP1 foci were not completely abolished by VE822 treatment, we noticed that the remaining foci were larger in size with respect to untreated cells, suggesting that they were not *de novo* recruited molecules but rather 53BP1 bodies persisting from unresolved damage during the previous cell division (Extended Data Fig. 4a)^50,51^. Of note, the ANP32E-dependent activation of ATR-mediated DDR was also confirmed in MDA-MB-231 and SUM-159-PT cell lines, showing a reduction of pRPA32 and 53BP1 recruitment upon ANP32E silencing and an increase upon ANP32E overexpression, respectively (Extended Data Fig. 4b-e).

**Figure 3.**
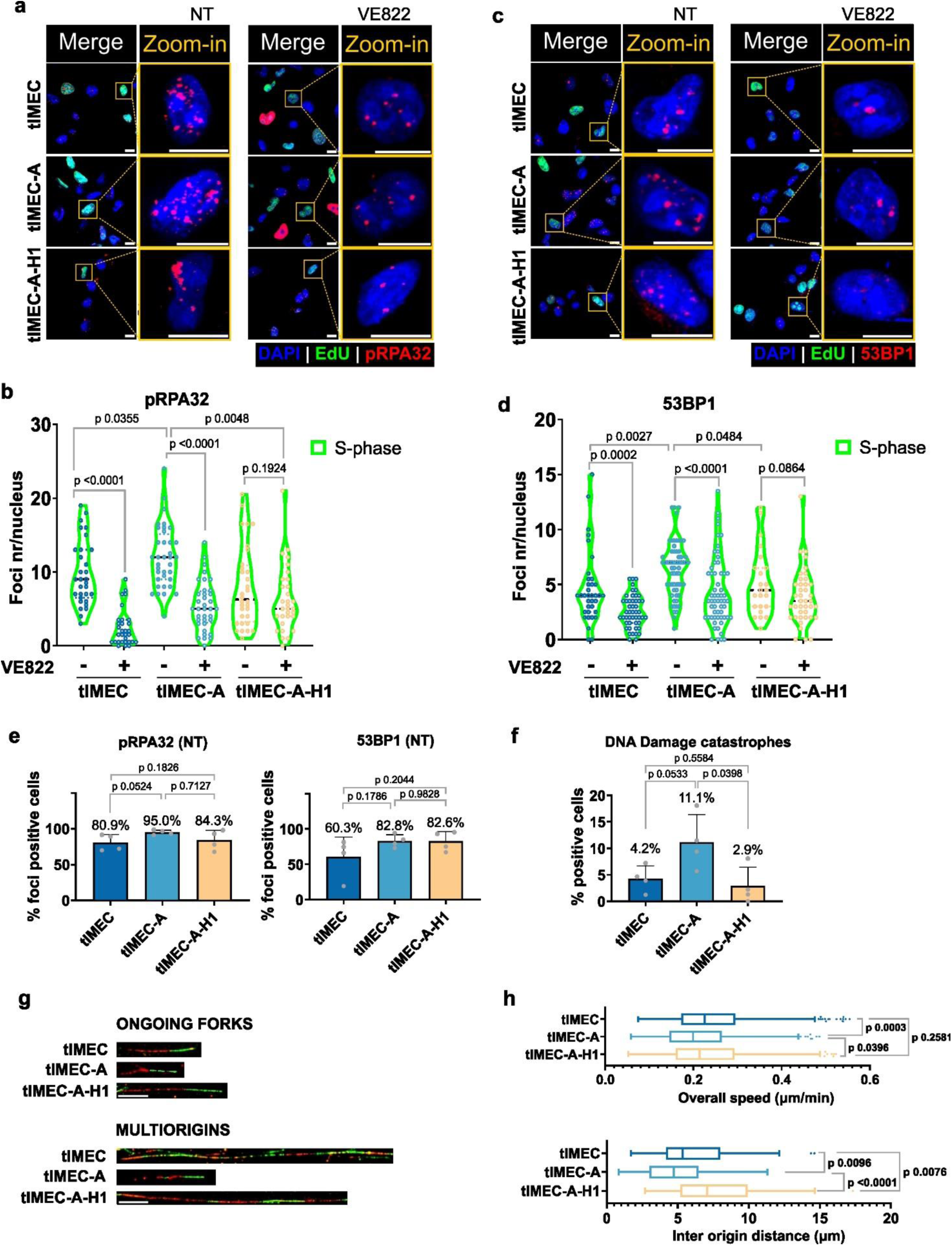
ATR-mediated DDR factors recruitment in actively replicating cells is ANP32E and R-loops dependent. (**a**) Representative images for p(S33)RPA32 immunofluorescence combined with EdU staining to label S-phase cells. Merge of DAPI (blue), EdU (green) and pRPA32 (red) is shown with zoom-in on EdU positive single cells where EdU channel has been excluded for clarity purposes. Non-treated (left) or VE822 0.25 µM 6h treatment (right) is shown. Scale bar = 10 µm. **(b)** Quantification of pRPA32 foci number per nucleus in EdU positive cells, with or without VE822 0.25 µM 6h treatment as indicated in the graph. P-values for unpaired t-test are reported above the compared plots. **(c)** Representative images for 53BP1 immunofluorescence combined with EdU staining to label S-phase cells. Merge of DAPI (blue), EdU (green) and 53BP1 (red) are shown with zoom-in on EdU positive single cells where EdU channel has been excluded for clarity purposes. Non-treated (left) or VE822 0.25 µM 6h treatment (right) is shown. Scale bar = 10 µm. **(d)** Quantification of 53BP1 foci number per nucleus in EdU positive cells, with or without VE822 0.25 µM 6h treatment as indicated in the graph. P-values for unpaired t-test are reported above the compared plots. **(e)** Barplots showing the percentage of foci positive cells for pRPA32 (left) and 53BP1 (right) in untreated condition. Mean and sd for 4 biological replicates is reported. P-values for unpaired t-test are reported above the compared plots. **(f)** Barplot showing the percentage of cells that upon VE822 0.25 µM 6h treatment show nuclei that are totally positive for pRPA32 staining. Mean and sd for 4 biological replicates is reported. P-values for unpaired t-test are reported above the compared plots. **(g)** Representative images of Ongoing replication forks (top) and Multiorigins replication forks (bottom). Scale bar = 10 µm. **(h)** Quantification of the overall replication fork origins speed (top) and inter-origin distance of multiorigins fibers (bottom) of a representative experiment. About 300 fibers were measured, of which 50-100 were multiorigin. P-values for unpaired t-test are reported above the compared samples.

DNA repair is a process that requires time to be accurately accomplished, at the expense of transcription and replication dynamics^52,53^. We therefore asked whether the increased frequency of TRCs and sustained DDR activation observed in ANP32E-overexpressing cells might have some drawbacks on transcription and replication speeds. We performed EU labeling of nascent transcripts after synchronizing transcription elongation with DRB treatment and washout^54^. Reduced levels of nascent RNA were already visible at the steady state in tIMEC-A cells, and the difference was accentuated upon transcription inhibition through DRB treatment and release of stopped RNApol II, revealing a slower transcriptional dynamic in tIMEC-A cells (Extended Data Fig. 4f, g). By performing a DNA fiber assay to measure replication fork speed^55^ we observed a slower progression of replication forks and an increased frequency of multiple origin fibers, with origins firing closer one to each other in tIMEC-A cells (Fig. 3g, h, and Extended Data Fig. 4h). R-loops resolution by RNaseH1 overexpression could restore the replication dynamics, suggesting a direct involvement of R-loops in impairing fork progression. In summary, ANP32E overexpression results in sustained activation of ATR-dependent DDR in actively replicating cells through a mechanism involving R-loops formation and altering transcription and replication dynamics.

### ANP32E overexpression leads to stalling of the elongating RNApol II

To determine the molecular mechanisms by which ANP32E elicits TRCs, we investigated whether its overexpression enhanced H2A.Z turnover genome-wide. By performing CUT&RUN, we found that ANP32E overexpression resulted in an increment of its chromatin binding at genomic sites enriched for H2A.Z and P400, a core component of the P400/TIP60 complex that interacts with ANP32E^24^ (Extended Data Fig. 5a). Of note, the overexpression of ANP32E reduced the levels of H2A.Z at open chromatin regions (Extended Data Fig. 5b, c), in line with previous findings^56^. On the same genomic site, both ANP32E and its partner P400 showed a reduction in the chromatin-bound fraction (Extended Data Fig. 5c). This is in line with the recent finding by which H2A.Z eviction by ANP32E originate stable ANP32E-H2A.Z dimers that remain trapped in the nucleoplasm^23,57^. We next employed CUT&RUN data for initiating (pS5) and elongating (pS2) RNApol II to evaluate ANP32E effect on RNApol II promoter-proximal pausing and its stalling along the gene body. By measuring the signal intensity along the gene body, we observed a decreased abundance of initiating RNApol II in the proximity of the TSS of ANP32E-overexpressing cells, possibly reflecting the less stable barrier posed by H2A.Z evicted at the +1 nucleosomes (Fig. 4a)^58,59^. This observation was further corroborated by analyzing the peak distributions for pS5 and pS2 RNApol II, which showed that in tIMEC the pausing sites of RNApol II is localized in the proximity of the TSS, while resulting in being placed further downstream in ANP3E-overexpressing cells (Fig. 4b). Further, we retrieved an increased distribution of pS2 RNApol II along the gene bodies of ANP32E-overexpressing cells (Fig. 4a, b), suggesting that elongating RNApol II molecules are subjected to nucleosome stalling events. To verify this finding, we calculated the Pausing Index (PI) or Stalling Index (SI), a measure of RNApol II promoter-proximal pausing or stalling along the gene body, respectively (Extended Data Fig. 5d). Quantitative evaluation of PI and SI revealed that, while initiating RNApol II showed a lower PI in ANP32E-overexpressing cells, the SI was significantly increased for the elongating RNApol II in tIMEC-A cells. This indicated that ANP32E overexpression is associated with more frequent stalling of RNApol II during transcription elongation. Interestingly, R-loops degradation rescued this effect, highlighting the key role of R-loops in the modulation of the correct RNApol II processivity (Fig. 4a, b and Extended Data Fig. 5d)^15^.

**Figure 4.**
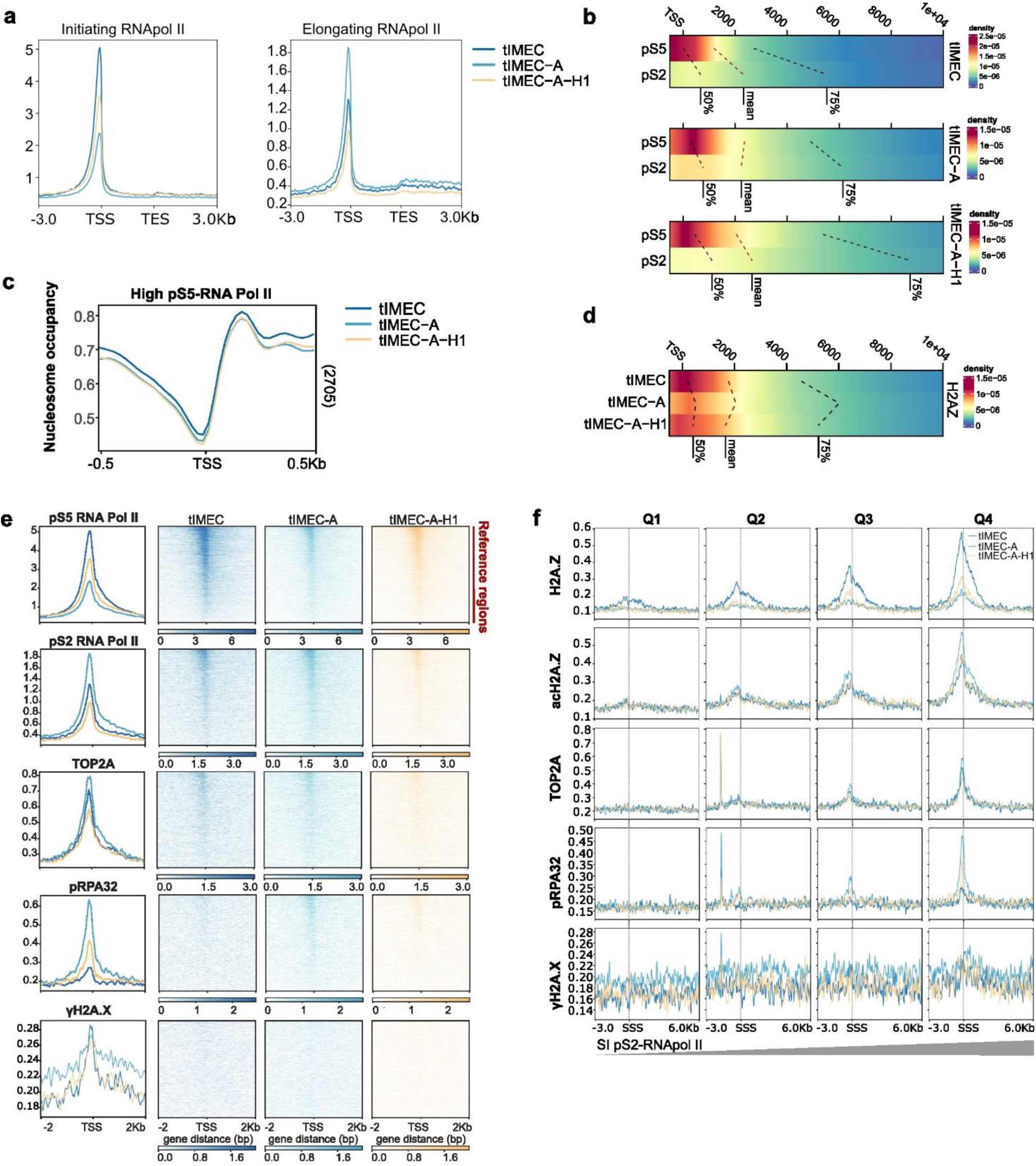
Determination of genome-wide chromatin effects of ANP32E overexpression. (**a**) Cumulative CUT&RUN signal plots of pS5– and pS2-RNApol II at pS5RNApol II-sorted ATAC-seq promoter clusters 1 to 5 (10,821 accessible sites). Regions were scaled on the gene body (TSS-TES) ±3 Kbp. **(b)** Heatmaps showing the density of pausing (pS5) and stalling (pS2) events, generated by plotting the distance from the nearest TSS of pS5– and pS2RNApol II peaks used in the calculation of PI and SI. 50%, mean and 75% of the peak’s distribution relative to TSS is reported in a window of –0.5 +10 kb from the TSS. Each of these values relates to a dashed line between pS5 and pS2 samples. **(c)** Nucleosome occupancy score cumulative plot at fourth (High level) quartiles of pS5RNApol II in tIMEC accessible regions. The number of considered genomic regions is indicated on the right of each plot. **(d)** Heatmpas showing the density of H2A.Z peaks at regions of highest pS2RNApol II stalling (fourth quartile), reported by plotting the distance from the nearest TSS. 50%, mean and 75% of the peak’s distribution relative to TSS is reported in a window of –0.5 +10 kb from the TSS and connected with a dashed line among the cell lines. **(e)** Cumulative CUT&RUN signal plots of acH2A.Z at RNApol II-sorted ATAC-seq promoter clusters 1 to 5 reported in Supplementary Figure S5B (10,821 accessible sites). Regions are centered on the TSS and a window of 500 bp up– and downstream are shown. **(f)** Cumulative plot of CUT&RUN immunoprecipitated factors on genomic regions of stalled pS2RNApol II grouped into quartiles according to the SI (n regions for every quartile = 2827). SSS indicates the Stalling Start Site. A window of –3Kb and + 6Kb from the PSS is displayed.

We hypothesized that the misregulated processivity of RNApol II directly depends on the altered H2A.Z turnover due to its effects on nucleosome stability. Therefore, we employed ATAC-seq data to calculate the nucleosome occupancy at sites enriched for initiating RNApol II and harboring H2A.Z (Extended Data Fig. 5e). In cells overexpressing ANP32E we observed a reduced occupancy of nucleosomes positioned downstream the TSS and within the gene bodies (Fig. 4c, d). Of note, this pattern was not preserved when we analyzed TSS characterized by a low abundance of initiating RNApol II (Extended Data Fig. 5f). By analyzing the peaks distribution for H2A.Z we found that while the relative abundance resulted diminished in tIMEC-A, it resulted unevenly distributed downstream the TSS (Fig. 4d). These findings suggest that the chaperone activity of ANP32E strengthened the nucleosome barriers at transcriptionally competent genes, resulting in stalling of the elongating RNA Pol II.

### ANP32E-dependent accumulation of DDR factors at RNA Pol II stalling sites

We hypothesized that the increased pausing of the RNAPol II may increase the frequency of TRC downstream of the TSS. Therefore, by using the initiating RNApol II as a reference of transcriptionally active genes (Extended Data Fig. 5g), we looked at the binding of TRCs-related DNA damage factors pRPA32 and γH2A.X, and TOP2A, which is a topoisomerase involved in the resolution of DNA supercoiling occurring in between the colliding transcriptional and replicative machinery^52^. We noticed that actively transcribed sites showed an increased recruitment of such factors in ANP32E-overexpressing cells, with pRPA32 being strongly enriched downstream of the TSS, while γH2A.X accumulated in the surrounding genomic regions (Fig. 4e). Of note, this pattern was partially rescued by the RNAsH1-mediated resolution of R-loops, suggesting a possible contribution in enhancing the loading of TRCs-related DNA damage factors (Fig. 4e). To clarify whether the TRCs-associated DDR factors are deposited in proximity of RNA Pol II stalling sites, we clustered the genomic regions based on the SI quartiles of elongating RNApol II and looked at their relative accumulation. Interestingly, we could first clearly observe the shift between pS5– and pS2-RNApol II from the start of the stalling site (SSS) with an intensity that correlates with higher SIs of elongating RNApol II (Extended Data Fig. 5h). This pattern was mirrored by a gradual increase of H2A.Z abundance in the surrounding of stalling sites, which was counteracted by the chaperone activity of ANP32E (Fig. 4f). Moreover, pRPA32 and TOP2A both accumulated at the stalling sites in ANP32E-overexpressing cells, while γH2A.X had a more widespread distribution, but mainly at or downstream of the PSS. Considering that an additional function of H2A.Z is to keep chromatin in an open state at sites of DSBs where its acetylation is necessary for the initial stages of DDR ^57,60,61^, we monitored the relative distribution of acH2A.Z (Fig. 4f). We found that regions with the highest stalling index accumulated acH2A.Z in an ANP32E-dependent manner, thus recapitulating the observed pattern of the TRCs-associated DDR factors. We excluded that the observed differences depended on the alteration of chromatin accessibility resulting from H2A.Z turnover since ATAC-seq signal increased proportionally with the magnitude of pausing but independently from ANP32E status (Extended Data Fig. 5i). Overall, our data indicate that the increased turnover of H2A.Z leads augmented stalling of the elongating complexes along the gene body, involving the formation of R-loops. The sites of increased pS2RNApol II stalling also show higher recruitment of DNA damage markers and retain acH2A.Z, indicative of active DDR.

### H2A.Z altered turnover primes for the formation of long R-loops

We next investigated whether R-loops accumulated at the ANP32E-associated damage sites by performing DRIP-seq. We found that ANP32E-overexpressing cells showed a higher number of R-loops peaks, whose abundance did not decrease upon RNaseH1 overexpression (Fig. 5a). We therefore looked at the length of R-loops reasoning that RNaseH1 degrades R-loops after they are formed, resulting in their shortening, but does not prevent their formation. R-loops resulted significantly longer in tIMEC-A cells with respect to both tIMEC and tIMEC-A-H1 (Fig. 5b, and Extended Data Fig. 6a). Interestingly, R-loop length increased at genomic sites that showed an H2A.Z loss in ANP32E-overexpressing cells (Fig. 5c, and Extended Data Fig. 6b). We observed a similar behavior at sites of reduced accessibility in tIMEC-A (Fig. 5d, and Extended Data Fig. 6c). Of importance, the expression of RNAseH1 re-established a lower R-loops length without affecting the relative abundance (Fig. 5c, d and Extended Data Fig. 6b, c).

**Figure 5.**
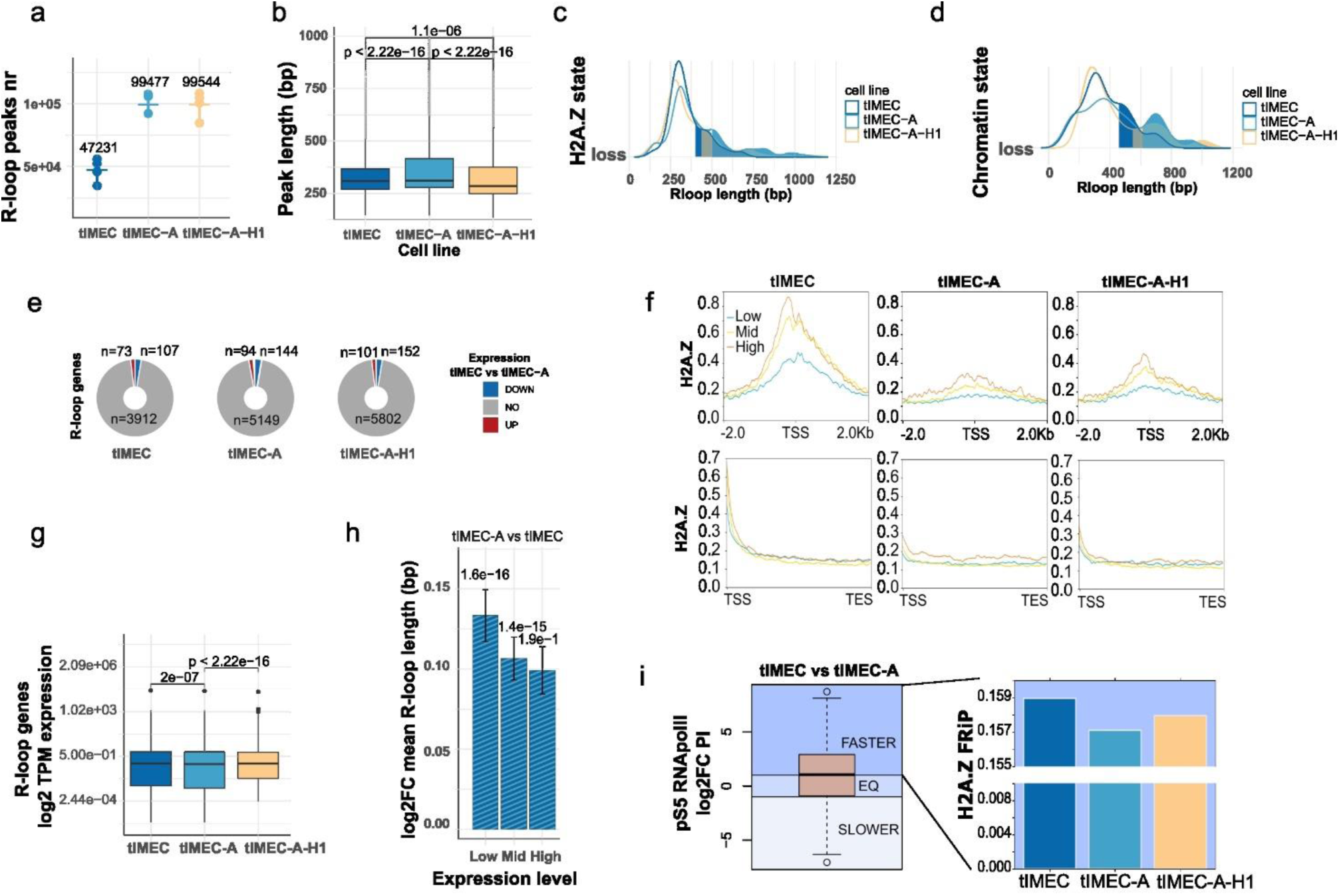
Longer and more frequent R-loops are formed in regions of H2A.Z eviction. (**a**) DRIP-seq peaks number calculated for the indicated cell lines, representing the abundance of R-loops. Peak number obtained for single replicates is shown with mean and sd. Mean peak number is reported above samples. **(b)** Boxplot reporting R-loop peaks length in bp in every cell line, as indicated on the x axis. P-values for unpaired t-test are reported above the compared plots. **(c)** Density plot representing R-loop peaks length in bp for R-loops intersecting regions of loss H2A.Z in tIMEC vs tIMEC-A. The 75° percentile is highlighted in the length distribution for every cell line. **(d)** Density plot representing R-loop peaks length in bp for R-loops intersecting regions of chromatin loss accessibility in tIMEC vs tIMEC-A. The 75° percentile is highlighted in the length distribution for every cell line. **(e)** Pie charts reporting R-loop genes number that have differential gene expression (UP or DOWN) or are unvaried (NO) for tIMEC vs tIMEC-A comparison as calculated from RNA-seq data. **(f)** Cumulative plots reporting the amount of H2A.Z deposition at the TSS (top) and gene body (bottom) of genes with High, Medium or Low expression, as established from RNA-seq data. **(g)** Expression values in log2TPM of tIMEC-A R-loop-forming genes. T-test p-values for the indicated comparisons are reported above graphs. **(h)** Barplot reporting the mean and sd of the log2FC of differential R-loop length for the comparison tIMEC vs tIMEC-A. R-loops were clustered in different groups based on the corresponding gene expression: Low, Mid, High. T-test p-value for statistical significance of the FC is reported above bars. **(i)** Left: boxplot showing the differential PI sites expressed as log2FCbetween tIMEC and tIMEC-A cells. Horizontal lines indicate the thresholds used to define region of Faster, Equivalent or Slower release of pS5RNApol II in tIMEC-A vs tIMEC. Right: Barplot for Fraction of Reads in Peak (FRiP) of H2A.Z in genomic regions where pS5RNApol II release is faster in tIMEC-A vs tIMEC.

To verify whether the enhanced R-loop formation in tIMEC-A was not solely a consequence of alterations in the transcriptome, we determined differential gene expression profiles by RNA-seq. This revealed that globally, about 500 genes have altered expression levels in tIMEC-A with respect to tIMEC cells, with a balance between up– and down-regulated genes, while less significant variation was observed upon RNaseH1 overexpression (Extended Data Fig. 6d, e). Of note, most of the genes harboring R-loop did not show a significant change in gene expression (Fig. 5e and Extended Data Fig. 6f), thus excluding a significant contribution of global transcriptional alteration to the enhanced formation of R-loops in ANP32E overexpressing cells. These findings strengthened the hypothesis that H2A.Z turnover is responsible for accumulating long R-loops. Considering that the distribution of H2A.Z occupancy along the gene is proportional to the transcriptional output^60^, we looked at H2A.Z status in relation to gene expression^62^ (Fig. 5f and Extended Data Fig. 6g). We found that low-expressed genes are characterized by a reduced level of H2A.Z at the promoter, yet it is relatively enriched along their gene bodies, in line with previous findings^62^. This trend is preserved among cell lines at promoters, but we observed a preferential depletion of H2A.Z in tIMEC-A cells at the gene body of low-expressed genes (Fig. 5f and Extended Data Fig. 6g). Of importance, R-loop-forming genes in tIMEC-A cells were, on average, less expressed (Fig. 5g). When looking at the differential length of R-loops between tIMEC and tIMEC-A cells, we observed that the genes with low expression levels were characterized by a more significant variation in R-loops length (Fig. 5h and Extended Data Fig. 6h). Thus, our data indicate that longer R-loops might be a consequence of the faster release of initiating RNApol II following the destabilization of the chromatin barrier due to H2A.Z removal. Indeed, the regions where H2A.Z is lost in tIMEC-A showed both longer R-loops and a faster release of the initiating RNApol II (Fig. 5c, i and, Extended Data Fig. 6i). Chromatin accessibility changes are instead less impactful on the areas of active pS5-RNApol II (Extended Data Fig. 6j). Altogether, we observed that ANP32E overexpression induces a global depletion of H2A.Z on chromatin, affecting particularly low-expressed genes. H2A.Z depletion is associated with a faster release of the initiating RNApol II, possibly leading to an increment of long R-loops.

### ANP32E overexpression induces R-loop dependent TRCs affecting genome stability

The consequence of R-loops formation is the exposure of the ssDNA strand opposite to that of the DNA:RNA hybrid. If not promptly resolved, this state may lead to DNA breaks and eventually to genomic fragility^15^. We thus hypothesized that the hyperactive DDR observed in ANP32E-overexpressing cells is linked to DNA damage occurring specifically at R-loops sites. To validate this observation at the genome-wide level, we intersected the genomic regions harboring R-loops with those characterized by DNA damage. Notably, we observed a more significant overlap between R-loops and the DDR factors pRPA32 and γH2A.X, as well as TOP2A, in ANP32E-overexpressing cells with respect to tIMEC, confirming that DNA damage occurs at R-loop sites (Fig. 6a). These results thus indicate that R-loops occur at sites of TRC and are in line with previous evidence showing that pRPA32 is recruited at the opposite ssDNA strand at TRCs sites^52,63^. By plotting the DRIP-seq signal at genomic regions with increasing pS2-RNApol II stalling, we noticed higher and more widespread R-loop occupancy in tIMEC-A cells, which was resolved upon expression of RNAseH1 (Fig. 6b). Given this correlation, and the substantialoverlap of R-loops with pRPA32 (Fig. 6a), we defined R-loop-dependent TRC sites as those genomic regions harboring both R-loops and pRPA32^63^ (thereafter named R-TRCs). Strikingly, we observed a very defined deposition of pRPA32 at the extremities of genomic regions characterized by long R-loops (Fig. 6c). Moreover, the elongating RNApol II occupancy matched with the visualized conflict site, indicating that we detected the transcriptional side of the TRC (Extended Data Fig. 7a). By inspecting the relative distribution of R-loops, pRPA32, and elongating RNApol II, we suggest that the conflict involved at least two transcriptional bubbles, as schematized above the graph (Fig. 6a Extended Data Fig. 7a). Interestingly, γH2A.X accumulated in between the two conflict bubbles, indicating the occurrence of DSBs (Extended Data Fig. 7a). Using R-loops and pRPA32 co-occupancy as a readout R-TRCs, we observed that the affected genes were more abundant in ANP32E-overexpressing cells, while genes with non-TRC R-loops (pRPA32-negative) or R-loop-independent sites of DNA damage (pRPA32 only) were equivalently shared independently from the cell line (Fig. 6d, Extended Data Fig. 7b). Consistently with what was described above, we also found that R-loops associated with R-TRCs were longer with respect to R-loops that did not lead to ATR-dependent DDR activation, especially in tIMEC-A cells, confirming that it is not only the higher frequency of R-loops formation but also the length of R-loops to correlate with DNA damage (Fig. 6e).

**Figure 6.**
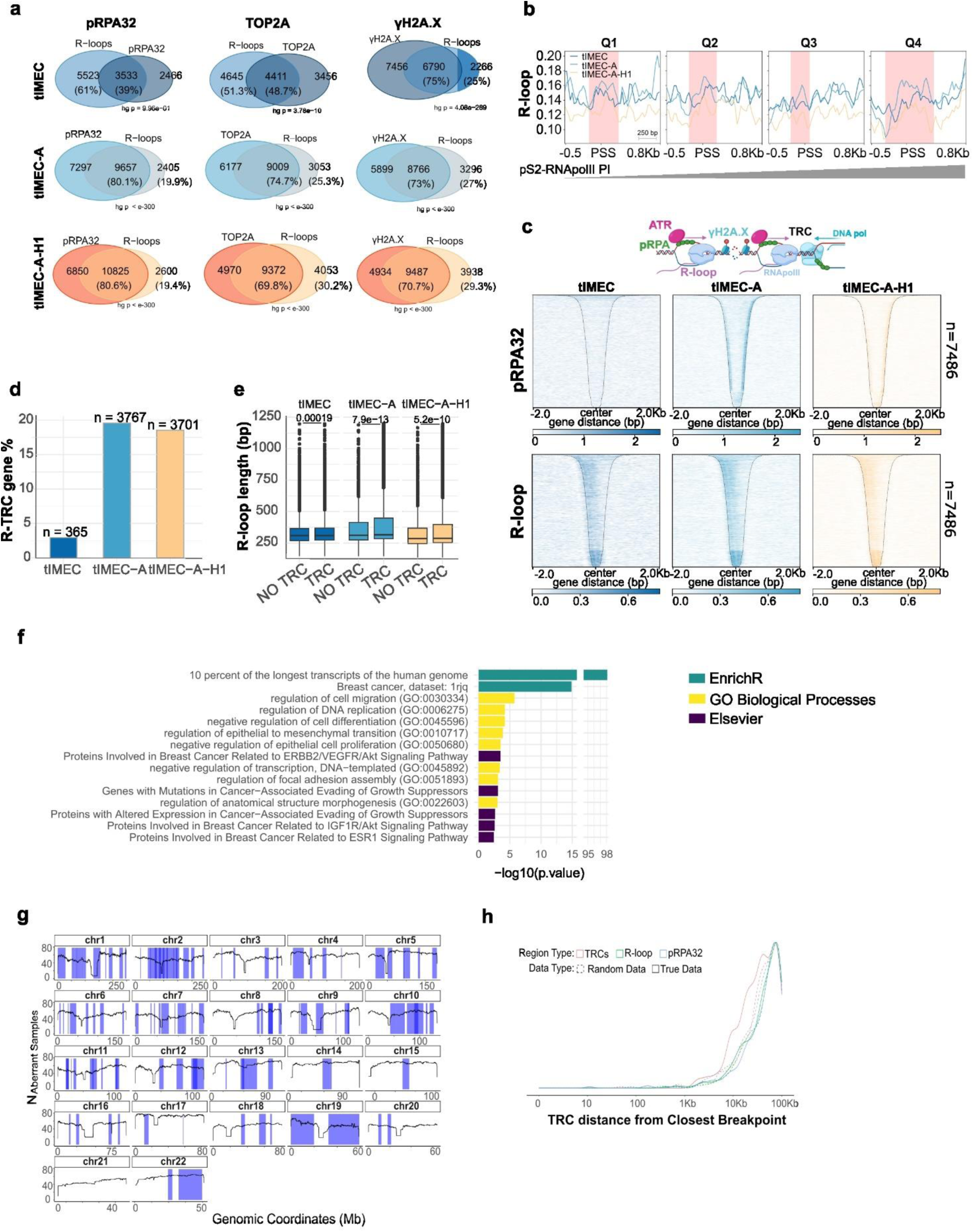
R-loops and DDR identify TRC sites and are a source of genome instability. (**a**) Intersection of genes that show R-loops and DDR markers according to DRIP-seq and CUT&RUN data, respectively. Hypergeometric test was used to evaluate statistical significance of the intersections. Raw gene number and percentage of overlapping genes are reported. **(b)** Cumulative plot showing R-loops distribution in respect to pS2RNApol II stalling sites, grouped into quartiles (Q1: lower stalling, Q4: higher stalling). Regions were centered on the stalling start site (PSS) and a window of –0.5 and +0.8 Kb is displayed. Red background indicates the extension of R-loops-covered region in tIMEC-A cells for every quartile of pS2RNApol II stalling. **(c)** Heatmaps showing the relative deposition of pRPA32 and R-loops at genomic regions occupied by both factors as derived from CUT&RUN and DRIP-seq, respectively. The displayed regions were ordered by decreasing size and the same order is maintained in each graph. The black dashes line indicates the end of the region. N regions = 7486. A scheme representing the proposed model of the TRCs is reported above the graph matching R-loops and pRPA32 deposition as in the tIMEC-A heatmap. **(d)** Barplot reporting the percentage of genes in which toxic TRCs (positive for pRPA32 and R-loops) were measured in each cell line. Raw number of genes is reported above each bar. **(e)** Boxplot of R-loops length in bp for TRC regions (pRPA32 + R-loop) and NO TRC (R-loop only) in the indicated cell lines. T-test p-value are reported on graph for the indicated comparisons. **(f)** Gene Ontology of genes characterized by toxic TRCs. Enriched terms with the respective –log10(p-value) are reported. Colors indicate the reference database from which the terms were found. **(g)** Plot indicating the CFSs span and genomic location (blue) for every human chromosome and the frequency of the alteration in Basal BC patients (black lines). **(h)** Density plot reporting the frequency of TRC regions (red) in relation to the distance from the closest CFS breakpoint. Regions characterized by non-toxic R-loops (R-loop only, green) or pRPA32 only (blue) are used as control regions as well as three sets of corresponding random regions (dashed lines).

Since ANP32E-overexpressing BC tumors correlated with an augmented aggressiveness and wort prognosis^29^, our next question was how R-TRCs could increase the fitness of cancer cells. Thus, we first performed a gene ontology to understand if the affected genes may have a role in BC. Strikingly, we found that several affected genes belong to pathways related to BC-associated processes and tumor progression, including deregulated proliferation and migration (Fig. 6f). Interestingly, about 80% of TRC genes in ANP32E-overexpressing cells are targets of ANP32E, reinforcing the direct involvement of ANP32E in the occurrence of TRC (Extended Data Fig. 7d). We thus investigated if R-TRCs affect genome stability. To do so, we retrieved Common Fragile Sites (CFS) that are associated with early stages of cancer development in patients and looked at the proximity between high fidelity TRC sites identified in our cell model and the closest CFS breakpoint (Fig. 6g)^64^. Notably, we found that TRC sites show higher proximity with CFSs with respect to regions with non-TRC R-loops and R-loops-independent sites of DNA damage (pRPA32 only); the robustness of this result was assessed by comparing the datasets against randomly generated regions (Fig. 6h). To strengthen the connection between R-loop-dependent TRCs and clinically relevant genomic instability, we stratified Basal BC patients from TCGA data based on the upregulation of FA pathway, which resulted in being up-regulated among patients harboring ANP32E amplification (Fig. 1d) and it is involved in the resolution of R-loops-dependent DNA damage^40^. Concordantly, we observed an enrichment of TRC at about 10kb from the closest Copy Number Variant (CNV), specifically in patients with active FA pathway (Extended Data Fig. 7e). To inspect if genomic instability affecting the identified regions may be connected to metastasis onset, we used public data and observed a higher frequency of genomic alterations in the actively expressed TRC-genes found in proximity to CFS breakpoints in patients with metastatic TNBC (Extended Data Fig. 7f). Together, these results indicate that R-loops-dependent TRCs occur at genes that belong to BC-related pathways in proximity to CFSs, possibly triggering genome instability.

### ANP32E overexpression triggers R-TRCs and vulnerability to ATRi in vivo

The obtained results indicate that ANP32E overexpression is associated with an increment of TRCs and DNA damage. To determine their pathogenic relevance in vivo, we orthotopically injected tIMEC and tIMEC-A cells in immunocompromised mice and characterized the formed tumors^35^.Tumor sections were analyzed by PLA to measure TRCs in vivo. We observed a strong enrichment in TRC foci in tIMEC-A-derived tumors, which was also reflected by an increase in ATR-dependent DNA damage measured through pRPA32 foci (Fig. 7a-c). These results are encouraging to propose ATRi treatment to trigger synthetic lethality in ANP32E-overexpressing tumors. To test this hypothesis, we first determined VE822 drug cytotoxicity in vitro, observing that ANP32E overexpression led to lower EC_50_ values in tIMEC-A and SUM-159-A TNBC models, reflecting increased vulnerability (Fig. 7d, Extended Data Fig. 8a, and Supplementary Table 2). Accordingly, ANP32E silencing in MB-231 cells, although it did not significantly affect the relative EC_50_ values, ameliorated cell growth and improved metabolic activity upon VE822 exposure (Extended Data Fig. 8b, c). Additionally, R-loops degradation by RNaseH1 overexpression in tIMEC-A partially rescued the drug cytotoxicity, supporting the key role of R-loops in triggering ATR-dependent DNA damage (Fig. 7d). The specificity of ATR inhibition as a synthetic lethal mechanism in ANP32E-overexpressing cells was also confirmed by using an alternative ATRi (Ceralasertib) and the ATR-independent DNA damaging agent ETP (Extended Data Fig. 8d, e). Caspase activation assay further confirmed ATRi efficacy on tIMEC-A cells, which displayed a significantly higher percentage of apoptotic cells compared to tIMEC and tIMEC-A-H1 (Extended Data Fig. 8f).

**Figure 7.**
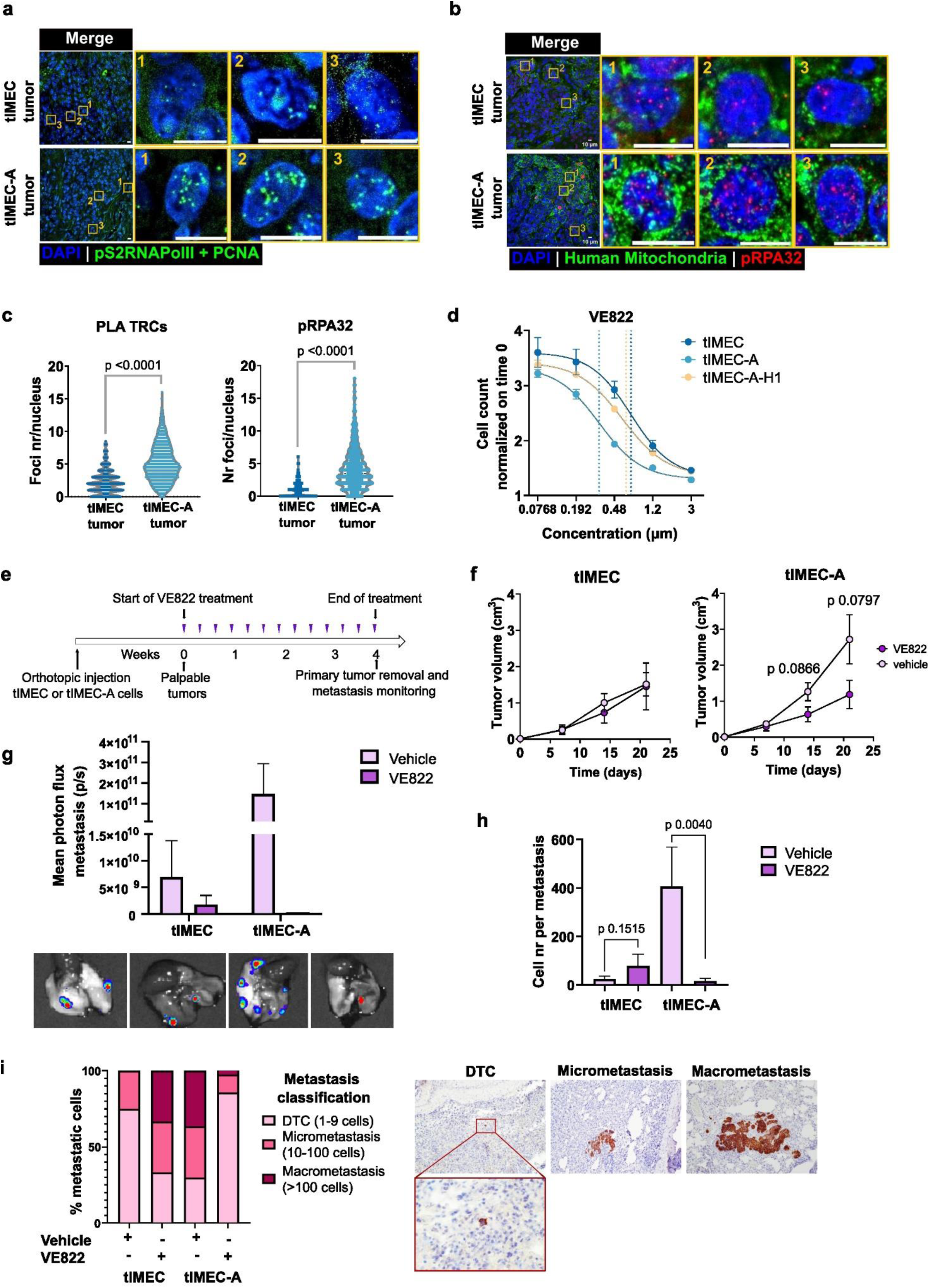
In vivo effect of ANP32E overexpression and ATRi treatment. (**a**) Representative images of TRC PLA between pS2RNApol II and PCNA obtained on tIMEC– and tIMEC-A-derived mouse xenografts. Three zoom-in panels of single cells are shown for both conditions. **(b)** Representative images of pRPA32 immunofluorescence (red) and human-mitochondria (green) obtained on tIMEC– and tIMEC-A-derived mouse xenografts. Three zoom-in panels of single cells are shown for both conditions. **(c)** Violin plots merging foci number/nucleus of two tIMEC– and tIMEC-A-derived xenografts per condition: TRC PLA (left), pRPA32 immunofluorescence (right), relative to representative images shown in (A) and (B), respectively. T-test p-values for the indicated comparisons are reported above graphs. **(d)** Representative drug response curves to 24h VE822 treatment of tIMEC, tIMEC-A and tIMEC-A-H1 cell lines. Vertical dotted lines indicate the respective EC_50_ value. **(e)** Scheme of the experiments used to test ANP32E overexpression in vivo and VE822 treatment effects. In brief, tIMEC or tIMEC-A cells expressing luciferase were orthotopically injected in mice and once the tumors became palpable, VE822 treatment started with four administrations per week. After 4 weeks tumors were resected, and metastases evaluated. **(f)** Dot plot showing the mean and s.e.m. of tumors derived from tIMEC (left) and tIMEC-A (right) cells in the presence or absence of VE822 treatment. Tumor volume was evaluated for 21 days when tumors became palpable. Two-way ANOVA p-values are reported for the significant comparisons VE822 vs vehicle. 4 to 5 mice per condition were analyzed. **(g)** Barplot showing the mean and s.e.m. photon flux per metastases of vehicle or VE822-treated mice harboring tIMEC and tIEMC-A-derived tumors. 2 to 3 mice per condition were analyzed. Images below graph report representative lung metastases as detected by luciferase bioluminescence. **(h)** Barplot showing the mean and sd for cell number per metastasis for tumors derived from tIMEC and tIMEC-A cells in the presence or absence of VE822 treatment. T-test p-values for the indicated comparisons are reported above graphs. **(i)** Left: stacked barplot of metastases classification based on number of cells found in every metastasis, as indicated in the figure legend. Right: Representative images of human-mithocondria IHC performed on tissue sections to identify DTC, micro– and macrometastases.

With these premises VE822 drug was administered in mice harboring tIMEC– and tIMEC– A-derived tumors (Fig. 7e). Mirroring patients’ data, ANP32E overexpression led to the formation of larger xenograft tumors, which resulted specifically responsive to ATR inhibition. Of importance, tIMEC-A-derived xenografts displayed a more aggressive phenotype characterized by large areas with higher proliferation index and poorly differentiated cellular elements bearing a high nucleus-to-cytoplasm ratio, representing a pathological feature of poor survival in cancer patients (Extended Data Fig. 8g, h). Conversely, tIMEC cells generated tumor xenografts with interspersed well-differentiated areas consisting of squamous epithelial cellular aggregates with modest eosinophilic cytoplasm and evidence of keratinization. Treatment with VE822 induced a more differentiated phenotype in tIMEC-A tumor xenografts, along with a reduction of ki67 expression (Extended Data Fig. 8g, h). Of importance, tIMEC-A-derived primary tumors resulted in being specifically responsive to ATR inhibition. Concomitantly to the formation of primary tumors, mice developed metastatic lesions, whose abundance was enriched in mice injected with ANP32E-overexpressing cells, yet this effect was strongly mitigated by ATR inhibition upon VE822 treatment (Fig. 7g)^29^. To better characterize the effect of ANP32E overexpression and ATRi on metastasis formation, we quantified the size of metastatic lesions. On average, metastases derived from ANP32E-overexpressed tumors were wider, and VE822 treatment strongly reduced their size (Fig. 7h). Specifically, ANP32E overexpression led to a higher frequency of micro– and macrometastases at the expense of DTCs. Importantly, VE822 treatment impaired the metastatic burden, resulting in the formation of scattered disseminated tumor cells (DTCs) that did not evolve into macrometastasis, at least at the time of observation (Fig. 7i). In summary, ANP32E overexpression is associated to increased TRCs and ATR-dependent DDR activation in vivo, uncovering a vulnerability to ATR inhibitor, which reduced the primary tumor volume and metastatic potential of BC cells.

## Discussion

The molecular processes leading to genomic instability in cancer cells are disparate, however RS has emerged as a common feature favoring its occurrence^2^. In the present work, we identified the overexpression of the histone chaperone ANP32E as a factor that exacerbates the MYC-induced RS, revealing a dependency of cancer cells on DNA repair pathways and, thus, a tumor vulnerability. We showed that ANP32E deregulation leads to an increased H2A.Z turnover, resulting in a higher frequency of RNApol II stalling, R-loop dependent TRCs, and genome stability.

It has been reported that genomic instability in cancer may derive from the deficiency in DNA repair genes. For instance, germline mutations in BRCA1/2 are found in 15 to 25% of all TNBC patients and are the only clinically validated biomarkers for personalized therapy^65^. Deficient DDR is not the case for patients with MYC-ANP32E overexpressing TNBC since we observed an upregulation of several of the DDR pathways concomitant with increased genomic instability. Indeed, the most upregulated DDR pathway is the FA, notoriously active in the resolution of R-loops-derived DNA damage in response to ATR signaling^39,40^. Concordantly, by recapitulating the deregulation of ANP32E in a basal BC model, we observed that its overexpression correlates with increased R-loops formation and FANCD2 ubiquitination which, thanks to ATR is recruited and retained at R-loop sites as part of FA pathway active cascade. In line with previous reports, we observed that concomitantly to R-loops increase there is an augmented frequency of Head-ON TRC. Importantly, Head-ON collisions are the most dangerous for DNA integrity and most often associated with the formation of R-loops and activation of ATR signaling^12^. Therefore, it is reasonable that ANP32E-dependent TRC may stimulate DNA damage, resulting in a sustained activation of DDR pathways, among which the FA pathway, recapitulating what was observed from patients’ data. Indeed, as confirmation of the DNA damage resulting downstream to ANP32E overexpression, we also measured an increase in the activation of the ATR-dependent DNA damage markers pRPA32 and 53BP1, which are indicative of DNA lesions. The described observations reflect the strong RS to which ANP32E-overexpressing cells are subjected and their dependence on ATR-mediated repair pathways to avoid genomic instability. As a proof of concept, treatment with an ATR inhibitor drug (VE822) triggered genomic instability in these cells, as reflected by the increase in DSBs, DSB-derived MNi, the occurrence of DNA damage catastrophes, and the accumulation of 53BP1 bodies^49,51^.

To unveil the possible molecular mechanism by which ANP32E contributes to genome instability, we determined its role in controlling transcription processivity and replication timing. We found that upon ANP32E overexpression, R-loop-dependent TRC facilitate the stalling of elongating RNApol II and its retention along the gene body and TTS. In this biological context, we also determined a spurious firing of replication origin, possibly contributing to the increased frequency of TRCs. Besides, R-loop dependent TRCs are known to impede the replicative fork restart at stalled sites, further exacerbating the firing of non-canonical origins^13,15^. Of note, H2A.Z is also enriched at replication origins in mammals and has a role in the regulation of early origins firing and replication timing^66^, indicating multiple layers of action that converge in the dysregulation of origin firings following ANP32E overexpression.

Furthermore, we unveil the contribution of H2A.Z turnover to this biological process by analyzing the alterations in its genomic distribution in ANP32E overexpressing cancer cells. We found a peculiar outcome of H2A.Z turnover, which is differential at promoters and gene bodies^62^. This evidence allowed us to propose a molecular mechanism by which ANP32E-mediated eviction of H2A.Z at promoter-proximal regions leads to a faster release of paused initiating RNApol II^21^, resulting in a downstream accumulation of elongating complexes and their stalling along the gene body. This is accompanied by an increased recruitment of DNA damage factors and the formation of long R-loops. Additionally, ANP32E preferentially evicts H2A.Z along the gene body of low-expressed genes, giving rise to the accumulation of long R-loops.

Based on previous studies, we mapped R-loop-dependent TRCs as those genomic regions harboring both R-loops and pRPA32 marks^63^. This allowed the genome-wide visualization of the transcriptional side of the conflicts. Given the relative distribution of R-loops, pRPA32, γH2A.X, and elongating RNApol II, we suggest that the conflicts involved at least two transcriptional bubbles, with frequent occurrence of DNA DSBs in between. Importantly, we found that the genomic regions that are affected by R-loops-dependent TRCs involve genes that give rise to long transcripts^8,13^. This result suggests a peculiarity of ANP32E action that specifically affects late-replicating, long transcripts, being responsible for the onset of CFS at these specific genomic sites. Importantly, TRCs were also enriched in genes of BC-associated pathways, primarily involved in cell migration and proliferation. According to the hypothesis that unresolved TRCs give rise to DSBs and ultimately to genome instability, we found that R-loops-dependent TRC formed consequently to ANP32E overexpression were significantly enriched in proximity of known CFSs. The biological relevance of this finding is further supported by recent results obtained from patients of breast adenocarcinoma relating APOBEC3-dependent kataegis in R-loop regions with similar proximity from structural variants breakpoints^67^.

Increasing the relevance of the molecular mechanism that links ANP32E overexpression to genome instability, we confirmed *in vivo* that ANP32E stimulates the recruitment of pRPA32 and the formation of Head-ON TRC. Moreover, we found that the inhibition of ATR with VE822 strongly reduced the formation of primary tumors and derived metastases. This result is of particular importance since it gives a rationale for proposing ANP32E expression status as a diagnostic marker, together with MYC amplification, to stratify patients that would benefit from ATRi-based therapies. Since the available clinical trials suggest that DDR inhibitor drugs, including ATRi, are more effective in combinational therapies, it would be worth evaluating possible drug combinations to maximize treatment success^68–70^. Moreover, a broader perspective can be given to this study by extending ATRi treatment to other types of cancers that overexpress ANP32E alone or in the presence of RS-inducing oncogenes different from MYC. In summary, here we proposed a molecular mechanism through which ANP32E overexpression can contribute to aggravate RS by inducing epigenetic changes that lead to the accumulation of R-loop-dependent TRC and genomic instability, which can be exploited for the rational administration of ATRi drugs to increment therapeutic success.

## Author Contributions

S.L., V.P. and A.Z. conceived the study, designed the experiments, and interpreted the data. S.L and L.F. performed chromatin, expression and R-loops profiling experiments and computational analyses. S.L, V.P., M.B., and A.F. performed the cellular and molecular biology studies, participating in data analyses. A.T. and M.G. performed the *in vivo* experiments and IHC analyses. M.T. designed the in vivo experiments and interpreted the data. Y.C., G.D’A. performed computational analyses of cancer patients’ datasets under the supervision of F.D. A.Z. supervised the work, S.L and A.Z. wrote the manuscript.

## Acknowledgments

We thank Maria Luce Negri and Dr. Sven Beyes, members of Zippo laboratory, for helpful discussion and technical support. We thank Dr. Veronica De Sanctis, Dr. Roberto Bertorelli and Dr. Paolo Cavallerio of the Next Generation Sequencing (NGS) core facility for their helpful technical contribution and sample sequencing. The Cell Technology (CTF) and High Throughput Screening (HTS) facilities at the CIBIO Department are also gratefully acknowledged. We thank Prof. Luca Fava, Prof. Alessandro Provenzani and Prof. Matthias Altmeyer for critical comments and suggestions. This project has been realized with the financial contribution from Associazione Italiana per la Ricerca sul Cancro (AIRC), project code 25373 (to S.L.). Funding from AIRC under IG 2018 – ID. 21492 project, – PNRR M4C2I1.3-HEAL ITALIA project PE 00000019 CUP B73C22001310006 from Italian Ministry for University – PRIN 2022 prot. 20223NY37M supported MT work. European Union-NextGenerationEU-Fondi MUR D.M. 737/2021-Project title: “Sviluppo di terapie innovative nel trattamento del carcinoma alla mammella” supported AT work. European Union-NextGenerationEU-Fondi MUR D.M. 737/2021-Project title: “Ruolo di MERTK nell’interazione tra cellule tumorali e il microambiente nella progressione del tumore alla mammella” supported MG work. Work in the Zippo group was supported by grants from the AIRC foundation (IG 2019-22911) and European Union under the Horizon 2020 Framework Programme H2020 Future and Emerging Technologies (801336; – PROCHIP).

## Methods

### Cell culture

All cell lines were cultured at 37°C and 5% CO_2_. tIMEC, tIMEC-A and tIMEC-A-H1 cells were maintained in DMEM-F12 (Gibco, 11320-084) supplemented with human EGF (DBA, AF-100-15-1mg), Insulin (Merck, I6634-250MG), hydrocortisone (Voden, 74144), and bovine pituitary extract (Gibco, 13028-014) and 1x PenStrep (Gibco, 15070-063). SUM159 cell line was cultured in Ham’s-F12 (Gibco, 11765054) supplemented with insulin 5 μg/ml (Merck, I6634-250MG), hydrocortisone 1 μg/ml (Voden, 74144) and Fetal Bovine Serum 5% (Gibco 10270106) and 1x PenStrep (Gibco, 15070-063). MDA-MB-231 cell line was cultured in DMEM (Gibco 11960044) supplemented with Fetal Bovine Serum 10% (Gibco 10270106), 1x glutamine (Gibco 25030024) and 1x PenStrep (Gibco, 15070-063).

### Generation of stable cell lines

3xFLAG-ANP32E cDNA was cloned in a pGK plasmid backbone (Addgene, 169744) to transduce tIMEC and SUM-159-PT cells and generate the tIMEC-A and SUM-159-A cell lines. SUM-159-A cell population was next seeded at single cells level for clonal selection of FLAG-ANP32E positive cells, which were screened by IF and WB. Three independent clones were next employed for experiment, named SUM-159-A^cl1,cl2,cl3^, respectively. tIMEC-A-H1 cell line was generated by stable nucleofection of ppyCAG_RNaseH1_WT plasmid (Addgene, 111906)^16^ using the P1 Primary Cell 4D-Nucleofector X kit (Lonza, V4XP-1032) and selected by keeping them in culture under hygromycin (Applichem, A5347) 500 ng/ml selection for at least two weeks. MB-231^sh1^ and MB-231^sh2^ cell lines are two different cell populations, generated by transduction of MDA-MB-231 (named in short MB-231) with lentiviral vectors carrying pLKO.1 plasmid (Addgene, 8453) cloned with shANP32E-808 (sh sequence: GGATTTGATCAGGAGGATAAT).

### Dot Blot assay

Genomic DNA was isolated from tIMEC, tIMEC-ANP32E and tIMEC-ANP32E-H1 starting from a 80% confluent 6-well plate. Cells were pelleted and washed in PBS 1x (Gibco, 10010023). Next the cell pellet was resuspended in Lysis Buffer (Tris-HCl pH 7.2 10 mM, EDTA 10 mM, NaCl 150 mM, SDS 0.4%) + Proteinase-K 4 mg/ml (Roche, 3115887001) and incubated for 2h at 55°C. Samples were next precipitated with 1V cold isopropanol, centrifuged for 2 min at 4°C max speed and washed with ice cold EtOH 70%. After centrifugation the pellet was dried at RT and resuspended in ddMilliQ water by incubation 1-2h at 42°C. Dot blot samples were prepared from 1 µg gDNA with or without RNaseH1 2500 U treatment (NEB, M0297) O/N 37°C.Samples were denatured in Denaturation Buffer (NaOH 400 mM, EDTA 10 mM in ddMilliQ water) for 5 min at 95°C and then kept on ice to avoid reannealing. Denaturation was stopped by addition of 1V Ammonium Acetate 2 M. After equilibration and washing of a nitrocellulose membrane 0.2 µm (GE Healthcare, 10600001) in SSC 10x buffer (NaCl 3M, Sodium citrate 0.3 M in ddMilliQ water, pH 7.0) the prepared samples were spotted on the membrane using a Dot Blot apparatus (Bio-Rad, 170-6545) and UV-crosslinked for 3 min at 1200 J/m^2^. Next the membrane was blocked with PBS-milk 5% and incubated with primary antibodies s9.6 1:500 (Millipore, MABE1095) and dsDNA 1:3000 (Abcam, ab273137) O/N 4°c. After washing, specie-specific HRP secondary antibody (Sigma-Aldrich, AP160P) 1:3000 was applied, and the membrane was imaged on a ChemiDoc apparatus (Biorad).

### Proximity Ligation Assay (PLA)

Cells were seeded on gelatin 0.1% (Sigma Aldrich, G1393) coated glass coverslips in a 24-well plate. After 24h cells were fixed with ice-cold MeOH for 10 min at –20°C, blocked and permeabilized in Blocking Solution (Goat serum 5%, BSA 5%, Triton 0.5% in PBS 1x) for 1h at RT. PLA was performed using Duolink in situ detection kit (Sigma-Aldrich, DUO92008) with Duolink PLUS anti-rabbit (Sigma-Aldrich, DUO92002) and MINUS anti-mouse probes (Sigma-Aldrich, DUO92004) according to manufacturer instruction. The following primary antibody were incubated for 2h at RT: anti-pS2RNApolII (Abcam, ab5095) + anti-PCNA (Merck, WH0005111M2), anti-pS2RNApolII (Abcam, ab5095) + anti-Rloop (Millipore, MABE1095), anti-ANP32E (LB-Bio, LS-C344600-400) + anti-Rloop (Millipore, MABE1095). DAPI 1:1000 (Invitrogen, D1306) diluted in PBS 1x was applied for 30 min at RT for nuclei staining. Images were acquired at Nikon Ti2 confocal microscope equipped with a 60x oil-immersion ocular objective. PLA foci quantification was performed using a custom-made ImageJ macro.

### Western Blot

Whole-cell lysates were extracted using RIPA lysis buffer (Tris-HCl pH 8.0 10 mM, EDTA 1 mM, NaCl 140 mM, SDS 0.1%) supplemented with protease inhibitor cocktail 1x (Roche, 11873580001), phosphatase inhibitors 1x (Sigma-Aldrich, P5726 and P0044), PMSF 1 mM (Thermo Fisher, 36978), Na3VO4 1 mM (Sigma-Aldrich, S6508), okadaic acid 1x (Sigma-Aldrich, 459616). Whole cell lysates were then subjected to sonication with a Bioruptor sonicator for 10 cycles of 30 sec on/ 30 sec off, followed by two centrifugation steps at max speed for 10 minutes each. Protein concentration was measured from supernatant using a BCA assay kit (ThermoFisher, 23250). Depending on the target protein, 15-35 μg of the lysate were run on SDS-PAGE gels and transferred using a iBlot2 apparatus (Invitrogen) on nitrocellulose membrane (Invitrogen, IB23001/2), then blocked in either PBST-milk 5% (anti-ANP32E Abcam ab5993, anti-FLAG Sigma Aldrich F1804, anti-V5 Cell-Signaling 13202; anti-H3 Cell-Signaling 13202 4499, anti-β-actin Sigma-Aldrich A5441) or PBST-BSA 5% (anti-H2A.Z Abcam ab4174). Primary antibodies were incubated at 4°C O/N. Membranes were imaged with ECL substrate (Cytiva, RPN2232) on a Bio-Rad Chemidoc imager.

### Alkaline Comet Assay

Comet assay was performed using Comet Assay Kit (Abcam, ab238544) according to manufacturer instructions. Briefly, 150,000 cells were seeded in 6-well plates, grown O/N and when indicated treated with either ETP 2.5 µM (MedChemExpress, HY-13629) or VE822 1.25 µM for tIMEC, 2.5 μM for MDA-MB-231, 4.5 μM for SUM-159 (MedChemExpress, HY-13902) for 24h. Cells were scraped, embedded in low melting agarose at 1/10 ratio (v/v) and distributed on a glass microscope slide. Cells were lysed by incubating the slides for 45 min in lysis buffer (NaCl, EDTA, DMSO, Comet lysis solution 10x provided with the kit, pH 10.0). After incubation in Alkaline solution (NaOH, EDTA) for 30 min at 4°C, cells were subjected to alkaline electrophoresis for 45 min at 300 mA. Slides were washed in ddH_2_O and ice cold 70% EtOH before letting the agarose dry at RT. Once dried the slides were incubated for 15 min at RT with Vista Green dye. Comets were acquired with FITC filter of a Leica DMIL LED epifluorescence microscope equipped with a Leica DFC 450C camera. About 150-300 cells for every condition were acquired and quantified using both OpenComet^71^ plugin for ImageJ and CometScore software^72^.

### Micronuclei (MN) Assay and centromere detection in MN

For MN assay 7000 cells/well were seeded in 96-well plates. After 24h 0.03 µg/ml Mitomycin C (Mito C, Sigma-Aldrich M5353), increasing concentrations of VE822 as indicated in figure legends, or fresh medium (NT condition) were and incubated at 37°C for 6h. Next Cytochalasin B (Sigma-Aldrich, C6762) 0.6 µg/ml was added and incubated for 30 h at 37°C to block cytokinesis. Cells were then washed with PBS 1x and fixed in 4% PFA for 10 min at RT. Nuclei and MNi were stained using Hoechst solution (Invitrogen, H3570) 1:2000 in PBS 1x upon 30 min incubation at RT. After washing with PBS 1x plates were acquired using an Operetta imager with 20x objective (Perkin Elmer). Three technical replicates per condition were acquired for a total of 1500-3000 BN cells per condition. The experiment was performed in three biological replicates. MNi in BN cells were quantified by adapting the parameters of Operetta MN quantification analysis function.

For the detection of centromeres in MNi of BN cells, 30,000 cells were seeded on gelatin 0.1% coated coverslips in 24-well plates, grown ON at 37°C and treated as described above. After fixation, cells were blocked and permeabilized in Blocking solution (BSA 1%, Goat serum 5%, Triton-X 100 0.5%, PBS1x) for 1h at RT. Centromeres were stained using CREST serum (Antibodies Inc, 15-234) diluted 1:250 in PBS-BSA 3% and incubated for 2h at RT. Cells were washed 3x in PBS 1x, then incubated for 1h at RT with a solution containing: secondary antibody Alexa anti-human 647 1:1000 Invitrogen, A21445, Phalloidin 555 (Abcam, ab176756) 1:2000 and Hoechst 1:2000 in PBS 1x. Images were acquired with a Leica SP8 confocal microscope using a 60x objective and 2x zoom. Phalloidin staining was used to identify MNi that are in a cytoplasm of BN cells, which divided after MitoC or VE822 treatment. An in-house ImageJ macro was used to measure CREST signal in MNi.

### Immunofluroescence and EdU staining for nascent DNA

For IF staining 35,000 cells were seeded on gelatin 0.1% coated coverslips in 24-well plates and grown ON at 37°C. EdU treatment was performed using the Click-IT EdU Imaging Kits (Invitrogen, C10337) by treating cells for 1h with EdU 7.5 µM. Next coverslips were washed in PBS 1x and cells fixed in cold MeOH for 10 min at –20°C. After PBS-BSA 3% washes, fixed cells were blocked and permeabilized in Blocking solution (BSA 1%, Goat serum 5%, Triton-X 100 0.5%, PBS1x) for 1h at RT. EdU detection was allowed by click-it reaction of Alexa-Fluor 400 as described in the Click-IT EdU Imaging Kits (Invitrogen, C10337). Primary antibodies against FANCD2 1:100 (NovusBio, NB100-182SS), pRPA32 1:1000 (Bethyl, A300-246A-8) and 53BP1 1:100 (Millipore, MAB3802) were diluted in Blocking solution and incubated at RT for 2h. Next, coverslips were washed in PBS 1x before incubation with Alexa-Fluor-647 specie-specific secondary antibodies (Thermo Fisher, A32728 and A-21245) and either Hoechst 1:2000 for EdU labelled cells or DAPI 1:1000 for simple IF. For simple IF all washing steps were performed in PBS 1x and primary antibody directly incubated after cells blocking and permeabilization. Images were acquired using either a Leica SP8 or Nikon Ti2 confocal microscopes. Imaging quantification was performed with custom-made ImageJ macros.

### EU labeling and DRB treatment

Cells were seeded on gelatin 0.1% coated coverslips in 24-well plates and grown ON at 37°C. Cells were treated with 100 µM DRB (Sigma-Aldrich, D1916) for 3h to inhibit transcription elongation, next medium containing DRB was washed out to allow for synchronized release of transcription. Nascent DNA was labelled through EU incorporation (Click-iT® RNA Imaging Kits, Invitrogen, cat. C10329) for 1h hour starting from either: 1h before end of DRB treatment, together with DRB release or 2h after DRB release. Next samples were fixed for EU staining in 4% PFA for 10 min RT, permeabilized in PBS-Triton 0.3% and stained according to manufacturer instructions. Nuclei staining was performed with DAPI 1:1000 incubated for 30 min at RT. Images were acquired using a Leica SP8 confocal microscope equipped with a 63x oil immersion ocular objective. Imaging quantification was performed with a custom-made ImageJ macro.

### DNA Fiber assay

DNA Fiber assay was adapted from^73^, briefly 200,000 cells were seeded in 6-well plates and grown ON at 37°C in order to reach about 50% confluency. CldU 25 µM (Sigma Aldrich, C6891) was added to the medium and incubated for precisely 20 min at 37°C. Next, IdU 250 µM (Sigma Aldrich, I7125) was added without intermediate washing and incubated for precisely 20 min at 37°C. Cells were washed twice in ice cold PBS 1x, trypsinized, pelleted by centrifugation at 200 g for 5 min RT and resuspended in ice cold PBS 1x at a density of about 400,000 cells/ml. 2 µl of the cell suspension were pipetted at the top a positively charged microscope slide (Epredia, J1800AMNZ), air dried for max 5 min and lysed for 2 min in 7 µl of Spreading Buffer (Tris-HCl pH 7.4 260 mM, EDTA 40 mM, SDS 0.5 % w/v). Slides were next tilted to an angle of 25°-40° until the cells droplet reached the bottom edge of the slide. Slides were air dried and fixed in 1 ml methanol:acetic acid 3:1 for 10 min RT, washed twice in ddH_2_O, rinsed and incubated in HCl 2.5 M for 1h at RT. After PBS 1x washes to remove the excess of HCl, the slides were blocked in PBS-BSA 1% + 0.001 % Tween-20 for 1h at RT and incubated with a mix of rat monoclonal anti-BrdU antibody (1:500, Abcam Ab6325) and mouse monoclonal anti-BrdU antibody (1:500, BD Bioscience 347580) for 1h at RT. After 3x washes in PBS 1x the slides were re-fixed in PFA 4% for 10 min RT, washed again in PBS 1x and blocking solution, and incubated for 2h at RT with secondary antibody mix of Alexa Fluor 647 anti-rat (Invitrogen, A-21247) and Alexa Fluor 488 anti-mouse (Invitrogen, 11001) 1:500 in blocking solution. Lastly, the slides were washed in PBS 1x, dried to remove the excess of liquid and mounted using aqueous mounting medium (Histo-Line, PMT030). Images were acquired using a Nikon-AX confocal microscope with 60x objective and 2048×2048 pxl resolution. Three biological replicates were performed and for each one about 300 fibers were analyzed using DNA Stranding software^74^.

### ATACseq samples and library preparation

ATAC-seq samples were prepared according to the Omni-ATAC protocol^75^. Three biological replicates were performed for tIMEC, tIMEC-ANP32E and tIMEC-ANP32E-H1 cells. For each sample, 100,000 cells were collected and centrifuged at 500 rcf 4°C for 5 min. Cells were then resuspended in 50 μL of cold ATAC-Resuspension buffer (RSB; Tris-HCl pH 7.4 10 mM, NaCl 10 mM, MgCl2 3 mM, distilled water) containing 0.1% NP40 and 0.1% Tween-20 and 0.01% Digitonin (Promega, G9441) and incubated for 3 min to allow cells permeabilization. Nuclei were next isolated by washing cells with 1 mL cold ATAC-RSB buffer containing 0.1% Tween-20, and centrifuged at 500 rcf 4°C for 10 min. Isolated nuclei were resuspended in 50 μL of Transposition mix (Illumina Tagment DNA Enzyme 100 nM, TD-Buffer 2X (kit from Illumina, 20034197), PBS 0.33%, 0.01% Digitonin, 0.1% Tween-20, distilled water) and incubated at 37°C for 30 min in a thermomixer with 1000 rpm mixing, in order to allow the transposition reaction. The transposed samples were purified using Zymo DNA Clean and Concentrator-5 Kit (Zymo Research, D4013) and PCR-amplified using customize primers (Supplementary Material) designed following Illumina Adapter Sequences manual (Illumina, Document #1000000002694 v16) instructions. PCR-enriched libraries were, then, purified using AMPure XP Beads (Beckmann Coulter, A63881). Libraries quality and size distribution was analyzed at 2100 Bioanalyzer Instrument (Agilent, Model G2939BA) using Agilent High Sensitivity DNA Kit (Agilent, 5067-4626). Samples were sequenced using an Illumina NovaSeq6000 instrument.

### CUT&RUN samples and library preparation

CUT&RUN protocol was adapted from ^76^: cells were seeded and grown for 24h at 37°C in order to reach 70-80% confluency and 250,000 cells per sample were collected afterwards. For TOP2A CUT&RUN, before collection cells were treated with Etoposide 5 μM for 1h at 37°C to improve TOP2A signal^77^. Cells were next centrifuged 3 minutes at 600 g, washed three times in 1.4 mL Wash Buffer (HEPES pH 7.5 20 mM, NaCl 150 mM, Spermidine 0.5 mM, cOmplete™ EDTA-free Protease Inhibitor Cocktail 100X (Merck, 11873580001), and distilled water) and resuspended in 1 mL Wash buffer. Meanwhile, 150 μl Concavalin-A slurry (Polyscience, 86057-3) per sample were activated by washing the bead slurry three times in 1.5 mL Binding Buffer (HEPES pH 7.5 20 mM, KCl 10 mM, CaCl2 1 mM, MnCl2 1 mM). Activated beads were added to the cells while gentle vortexing and left rotating (25 rpm) for 10 minutes at RT. Afterwards, cells bound to beads were divided into aliquots, one for each primary antibody to be used (Supplementary Material). The supernatant was removed by placing samples on a magnetic stand, the beads were resuspended in 100 μL of Antibody buffer (Wash buffer, 0.025% Digitonin (Merck, 300410), EDTA 2 mM) and 1 μg of the antibody was added. Tubes were left rotating ON at 4°C. Samples were washed twice in 1 mL Dig-Wash buffer (Wash buffer, 0.025% Digitonin) and resuspended in 150 μL pAG-MNase 700 ng/ml diluted in Dig-Wash Buffer while gentle vortexing; tubes were left rotating (25 rpm) for 1h at 4°C. Samples were washed, resuspended in 100 μL DW Buffer and put on ice, next 2 μL of CaCl2 100 mM was added to allow chromatin digestion reaction. After 30 minutes of incubation 100 μL of 2X STOP buffer (NaCl 340 mM, EDTA 20 mM, EGTA 4 mM, 5% Digitonin, 100 μg/mL RNase A (Merck, R6513), 50 μg/mL Glycogen (Thermo Scientific, R0561), and distilled water) were added and incubated 30 min at 37°C. The supernatant containing digested chromatin was transferred to clean 1.5 mL DNA LoBind® tubes. Samples were phenol/chloroform extracted and quantified using a Qubit™ 4 Fluorometer (Invitrogen, Q33238) using the Qubit™ 1X dsDNA High Sensitivity kit (Invitrogen, Q33265). 3 ng of DNA were used to perform library preparation with NEBNext® Ultra™ II DNA Library Prep Kit for Illumina® (E7645), following the modified protocol from^78^ specific for small-fragment libraries. Libraries quality and size distribution was analyzed at 2100 Bioanalyzer Instrument (Agilent, Model G2939BA) using Agilent High Sensitivity DNA Kit (Agilent, 5067-4626). Samples were sequenced using an Illumina NovaSeq6000 instrument.

### Pausing Index analysis

Pausing index (PI) is determined by the ratio of read density at the promoter to the read density at the corresponding gene body^79^; therefore, the PI for pS5– and pS2-RNA Pol II was determined to evaluate the differential presence of stalled elongating enzyme along the gene body of Pol II occupied genes. The PI analysis was performed giving pS5– and pS2-RNA Pol II BigWig files to the getPausingIndices() function of BRGenomics (Version v1.8.0; DeBerardine M., 2022; https://bioconductor.org/packages/release/bioc/html/BRGenomics.html) R package (4.1.2.2 R software). PI was calculated at common pS5– and pS2-RNA Pol II annotated genes respectively, longer than 1 kb, in all three cell lines, defining the promoter region as –50/+300 bp interval from the TSS and the gene body as +301 bp from the TSS and 1 kb downstream the TES.

### Nucleosome occupancy calculation

NucleoATAC tool was employed for nucleosome mapping analysis starting from ATAC-seq data^80^. The analysis was performed on biological replicates merged reads, with genomic regions set as ±500 bp windows from line specific accessible TSS. Plots where produced using deepTools (4.1.2.3 DeepTools) computeMatrix and plotProfile utilities, using as input nucleoatac_signal.smooth.bedgraph.gz or occ.bedgraph.gz files converted to BigWiG.

### DNA:RNA immunoprecipitation and sequencing (DRIPseq)

DRIPseq samples were prepared based on the protocol described in Sanz et al. 2019^81^. Briefly, genomic DNA was extracted in native conditions and digested by restriction enzyme cocktail fragmentation (BsrGI-HF NEB cat. R3575, EcoRI-HF NEB cat. R3101, HindIII-HF NEB cat. R31045, SspI-HF Neb cat. R3132 and XbaI NEB cat. R01455). Immunoprecipitation was performed using 8 µg of fragmented DNA incubated for 14h at 16°C with 20 µg of S9.6 antibody (Millipore, cat. MABE1095). Next the complexes were captured on ProteinA/G agarose beads (ThermoFisher Scientific R0561), washed, eluted and purified by phenol/chloroform extraction. qPCR of control regions was performed to check the quality of the material. Libraries for sequencing were prepared by using NEB-Next end repair module (NEB, cat. E6050), followed by Klenow fragment exo-A-tailing (NEB, cat. M0212), adaptors ligation with Quick Ligation kit (NEB, cat. M2200) and amplification with Phusion Flash HF PCR master mix (ThermoFisher Scientific, cat. F548S). All purification and size selection steps were performed using AMPure XP Beads. Libraries were sequenced on an Illumina NovaSeq6000 platform with SR 100 bp in order to obtain 40-60Mreads per sample.

### RNA-seq

Total RNA for RNA-seq experiments was extracted from 75% confluent cells. Then, 1 million cells were trypsinized, pelleted, and resuspended in 1 ml TRIzol (Invitrogen, 15596018). Total RNA was extracted by phenol/chloroform and purified through the RNA Clean and Concentrator-25 kit (Zymo Research, R1018), following the manufacturer’s instructions. RNA integrity was checked through Agilent Bioanalyzer on Agilent RNA 6000 Pico Chips (Agilent technologies, 5067-1513). 450 ng of total RNA were next subjected to library preparation using the Universal Plus Total RNA-seq kit (TECAN, 9156-A01) according to manufacturer instructions. Libraries were purified using using AMPure XP Beads and fragment distribution checked through Agilent DNA High Sensitivity Chips (Agilent Technologies, 5067-4626). Samples were sequenced on an Illumina NovaSeq6000 platform with SR 100 bp in order to obtain 40-60Mreads per sample The experiment was done in four independent biological replicates.

### Data analysis

Raw FASTQ reads of CUT&RUN, ATAC-seq and DRP-seq were trimmed using Trimmomatic tool (https://chipster.csc.fi/manual/trimmomatic.html) to remove adaptor contamination and aligned to the primary assembly of the human reference genome version GRCh38 using Bowtie2 (https://bowtie-bio.sourceforge.net/bowtie2/manual.shtml) with –very-sensitive and – dovetail options for CUT&RUN data. CUT&RUN bam files were downsampled using BEDOPS bedextract utility (https://bedops.readthedocs.io/en/latest/index.html) to obtain the same reads number for every samples and assess unbiased comparison. ATAC-seq and DRIP-seq were instead normalized on the fraction of reads in peaks and library size respectively by using bedtools (https://bedtools.readthedocs.io/en/latest/). To account only for specific signal, DRIP-seq the ratio over RNaseH1 exogenous treatment was performed from bigwig files and only the regions with positive enrichment over noise were kept. Peak calling was performed with MACS2 in the case of CUT&RUN and DRIP-seq data (https://github.com/jsh58/MACS) by using –broadpeak option and a p-value of 0.00001 and 0.0001, respectively, and with Genrich tool (https://github.com/jsh58/Genrich) using a p-value of 0.001, AUC 100 and maximum peaks distance of 50 bp, PCR duplicates and ChrM corresponding reads were removed. ATAC-seq differential analysis was performed using DiffBind R package (http://bioconductor.org/packages/release/bioc/html/DiffBind.html). Peaks annotation was performed with Homer software annotatePeaks.pl function (http://homer.ucsd.edu/homer/) and the frequency of each annotation was normalized according to their overall genomic abundance in bp retrieved from UCSC Table Browser (https://genome.ucsc.edu/cgi-bin/hgTables). deepTools module CountReadsPerBin was used to calculate Fraction of reads in peaks (FRiP) by sampling the genome into 10000 positions of size 1 bp (https://deeptools.readthedocs.io/en/develop/source/deeptools.html). RNA-seq reads were aligned to the human reference genome with STAR (https://github.com/alexdobin/STAR/blob/master/doc/STARmanual.pdf) generating gene counts and removing PCR duplicates. Bam files were normalized according to the library size using bedtools. Transcripts expression levels were quantified as transcripts per million by mean of Salmon (https://salmon.readthedocs.io/en/latest/salmon.html#using-salmon). PATCHED chromosomes and mtDNA were removed from the analysis and differentially expressed genes were identified using the Bioconductor package DESeq2 and “apeglm” for LFC shrinkage (https://bioconductor.org/packages/devel/bioc/manuals/DESeq2/man/DESeq2.pdf). GO terms enrichment analysis was performed using EnrichR (https://cran.r-project.org/web/packages/enrichR/vignettes/enrichR.html).

All the further statistical analysis was performed using R software.

### Citotoxicity curves and EC50 calculation

To evaluate the cytotoxicity of VE822, Ceralasertib (MedChemExpress, HY19323) and Etoposide, cells were plated in 96-well plates, grown O/N at 37°C and treated with serial dilutions of the indicated compounds concentrations. Cell numbers were counted before treatment and 24h after treatment using Incucyte® Live-Cell Analysis System (Sartorius) that automatically counts cells based on dedicated analysis adapted on specific cell lines morphology. EC_50_ values were calculated based on the dose response curve fitting with a dedicated analysis available from Incucyte 2021C software. The dose-response curve was generated using the ratio of cell counts per image normalized on the counts at time 0h. 5 images per each well were acquired and each condition was prepared at least in technical duplicate.

### Caspase assay

To evaluate early apoptosis in cells treated with VE822, CellEvent Caspase-3/7 Green ReadyProbes™ Reagent (Invitrogen, R37111) was used. In brief, cells were plated in 96-well plates, grown O/N at 37°C and treated with different dilutions of VE822 together with the CellEvent™ Caspase-3/7 Green reagent consisting of a DEVD peptide. Cell growth was monitored for 48h after treatment using Incucyte® Live-Cell Analysis System (Sartorius) equipped with a FITC filter allowing the detection of cleaved caspase 3/7 leading to the cleavage of DEVD peptide and the release of green fluorescence. The percentage of apoptotic cells was calculated using the Incucyte® Live-Cell Analysis System 2021C software.

### Cell titer blue

To evaluate the cytotoxicity of VE822, cells were plated in 96-well plates at least in technical duplicate for each condition to be tested, grown O/N at 37°C and treated with serial dilutions of the indicated compounds concentration. At the end of treatment cells were incubated O/N with CellTiterBlue reagent (Promega, cat. G8080). The day after, 48h after treatment, fluorescence was acquired on a Varioskan plate reader (ThermoScientific). Fluorescence values were converted in percentage of viable cells based on the non treated control. Technical replicates were used to calculate mean and sd. Each experiment was performed at least in biological triplicate.

### In vivo tumor analysis

200,000 tIMEC or tIMEC-A cells, transduced with a lentiviral vector encoding for luciferase in 1:6 matrigel (BD Biosciences) were orthotopically injected in 5-week-old NSG mice (Charles River Laboratories). Once tumor xenografts were palpable (0.2×0.2 cm), mice were randomized (n=6) and treated with vehicle (10% D-a-tocopherol polyethylene glycol 1000 succinate in PBS, Sigma Aldrich) or VE822 (60 mg/kg) administered by oral gavage every other day for 4 weeks. Tumor volume was measured two times a week with an electronic caliper using the formula π/6 x (smaller diameter)2 × larger diameter. At the end of the treatments, bioluminescent signals were measured by using IVIS Lumina III system (PerkinElmer) and, after mice sacrifice, primary tumors and lungs were collected for immunochemical analyses.

### FFPE tissue immunofluorescence and PLA

For immunofluorescence and PLA experiments 5-μm tick FFPE sections of primary tumors were subjected to antigen retrieval by performing sections rehydration with successive incubations in xylene, ethanol 96%, ethanol 70%, ddH_2_O followed by 20 minute incubation in sodium citrate solution pH 6.0 10 mM or Tris-EDTA pH 9.0, respectively both supplemented with Tween-20 0.05% at 90°C. Next sections were blocked in AB solution (Goat serum 3%, Triton 0.3% in PBS) for 1h RT and incubated overnight with primary antibodies against pRPA32 1:1000 (Bethyl, A300-246A-8), anti-human-mitochondria (Abcam, ab92824) 1:1000 or anti-pS2RNApolII (Abcam, ab5095) + anti-PCNA (Merck, WH0005111M2) 1:400. Samples were washed in PBS-triton 0.3% and either incubated with specie-specific secondary antibodies Alexa-Fluor-647 and Alexa-Fluor-488, or subjected to PLA using Duolink in situ detection kit (Sigma-Aldrich, DUO92008) with Duolink PLUS anti-rabbit (Sigma-Aldrich, DUO92002) and MINUS anti-mouse probes (Sigma-Aldrich, DUO92004) according to manufacturer instruction.

### FFEPE tissue immunohistochemistry

Immunohistochemical analyses were performed on 5-μm tick FFPE sections of primary tumors and lung metastases as previously described^35^. Sections were exposed to ki-67 (D3B5, Cell Signalling Technology) or mitochondria (113-1, Abcam) antibodies. The number of lung metastatic cells was assessed by counting human-mitochondria positive cells with an optical microscope. For Hematoxylin/Eosin, tissues were stained with hematoxylin for 2 min and subsequently with eosin for 1 min.

### Data availability statement

Public BC patients data were retrieved from cBioPortal (https://www.cbioportal.org) using the brca_tcga_pan_can_atlas_2018 dataset (Breast Invasive Carcinoma TCGA PanCancer Atlas) and Breast Cancer METABRIC^27,28^ for Breast Cancer Specific analysis and the Metastatic Breast Cancer Project 2021 dataset (www.mbcproject.org, Provisional, December 2021), for metastatic BC patients analysis. Chromatin player genes were retrieved from GO terms: GO:0042393 and GO:0140713. The list of DDR pathway genes used for expression correlation analysis with ANP32E, MYC and FOXA1 was retrieved from ^32^. ANP32E mRNA expression data in BC cell lines were retrieved from the public dataset 23Q2 available from DepMap portal (https://depmap.org/portal/). All sequencing data produced in the present manuscript were deposited on GEO portal under accession nr GSE 250562.

