## Supplementary Data for "ANP32E drives vulnerability to ATR inhibitors by inducing R-loops-dependent Transcription Replication Conflicts in Triple Negative Breast Cancer"

### Extended Data

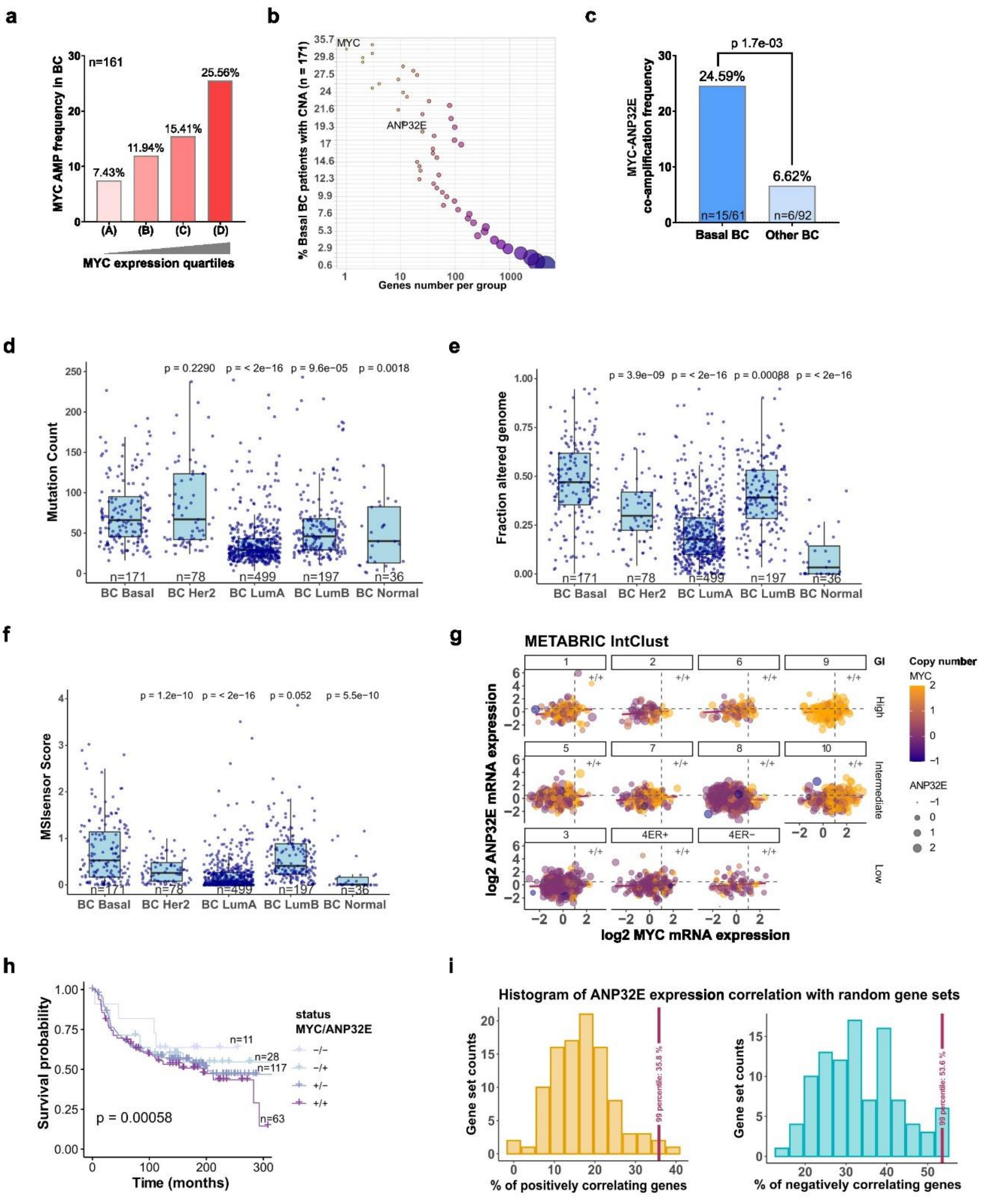


Lago_Extended Data Figure 1

**Extended Data Fig. 1. ANP32E is the most upregulated chromatin remodeler in Basal BC according to TCGA data.**

**(a)** Frequency percentage of BC patients with genomic CNA of MYC gene in relation to MYC mRNA expression z-scores relative to normal samples (log RNA Seq V2 RSEM) retrieved from TCGA data. Patients were grouped according to MYC mRNA expression quartiles, n=161. Quartiles: (A) -6.40 - -2.53, (B) -2.53 - -1.43, (C) -1.42 – 0.53, (D) -0.53 – 2.80. Enrichment in quartile (D) p-value= 4.11e-08. Data retrieved from TCGA. **(b)** Plot showing the frequency of CNA of specific genes in Basal BC patients. The number of genes with the same frequency is indicated on the x-axis and is proportional to the dots size in the graph. MYC and ANP32E genes are highlighted in the plot. Data retrieved from TCGA. **(c)** Frequency percentage of ANP32E CNA in patients with genomic CNA of MYC gene and Basal-BC or Other-BC subtypes. Data retrieved from TCGA. **(d)** Boxplot showing the Mutation Count of BC patients, grouped according to the BC subtype. Statistical significance was measured using t-test. **(e)** Boxplot showing the Fraction of Altered genome of BC patients, grouped according to the BC subtype. Statistical significance was measured using t-test. **(f)** Boxplot showing the MSIsensor score of BC patients, grouped according to the BC subtype. Statistical significance was measured using t-test. **(g)** Scatterplot showing the correlation between ANP32E and MYC mRNA expression (log2 z-score relative to all samples) in patients grouped as IntClust as defined for the METABRIC dataset. IntClust are sorted based on the genomic instability (GI) reported level (Low, Intermediate, High). Dot size is proportional to ANP32E gene copy number, while color scale blue to yellow reflects MYC copy number. The upper right quadrant indicates patients that are double positive for ANP32E and MYC overexpression. Dashed lines indicate the 75° of ANP32E and MYC expression, respectively. **(h)** Kaplan-Meier plot representing the survival probability of Basal BC patients with the different combinations of ANP32E and MYC upregulation. Data retrieved from Metabric dataset. Double positivity was assessed based on 75° of expression for both genes and concomitant copy number gain. P-values indicate statistical significance for the comparison of groups calculated by log-rank test. **(i)** Histogram showing the distribution percentage of 100 positively (left) or negatively (right) correlating random gene sets with ANP32E expression. The correlation score was calculated with Pearson method, considering a threshold of +/- 0.85. The sample size of random gene sets was established based on the mean gene nr of the DDR pathways (gene sets size = 28).

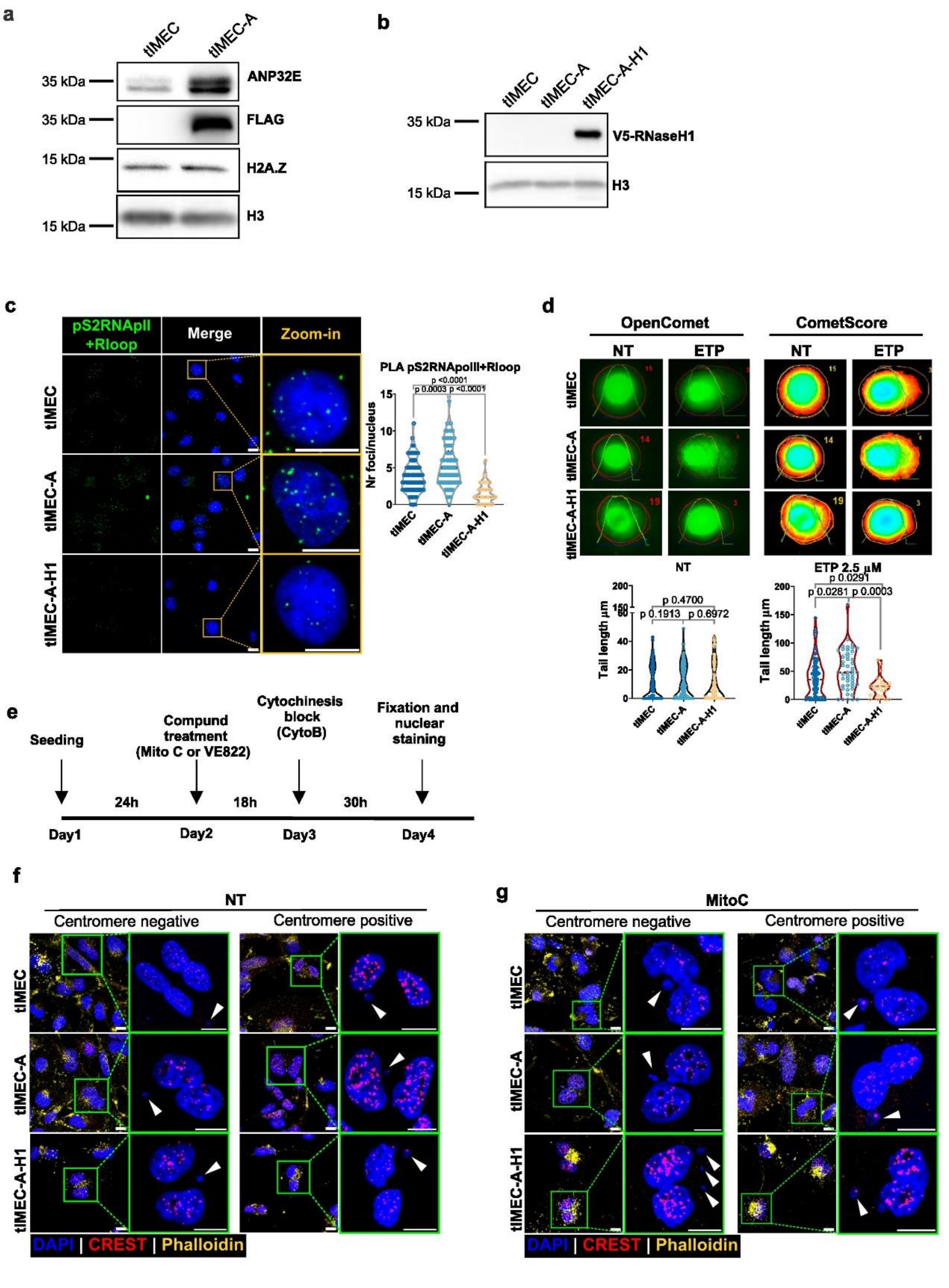


Lago_Extended Data Figure 2

**Extended Data Fig. 2. Establishment of ANP32E overexpressing basal BC model and effect on TRCs and DNA damage.**

**(a)** WB showing 3xFLAG-ANP32E overexpression and H2A.Z in tIMEC and tIMEC-A cells. Histone H3 is used as internal normalizer. **(b)** WB showing V5-RNaseH1 overexpression in tIMEC-A cells, used to generate tIMEC-A-H1 cell line. Histone H3 is used as internal normalizer. **(c)** Representative PLA foci images (left) and quantification (right) representing proximity between pS2RNApolII and R-loops. tIMEC n=80, tIMEC-A n=70, tIMEC-A-H1 n=47. Unpaired t-test p-values are reported in the graph. Scale bar = 10 µm. **(d)** Alkaline comet assay performed in the presence or absence of 24h ETP 2.5 µM treatment. Upper panel shows representative comets with quantification metrics for comet head and tail calculated by OpenComet ImageJ plugin[1] and heatmaps showing the DNA staining intensity measured by CometScore software[2]. The lower panel shows violin plots of comet tail length quantification. 150-300 cells were analyzed for every condition in 3 biological replicates. Unpaired t-test was applied to calculate p-value of statistical significance as reported in graph. **(e)** Scheme summarizing MN assay treatments and timings: cells were seeded and kept in culture for 24h, next either Mitomycin C, VE822 or no drugs were added to the medium. After 18h cytochalasin B was added to block cytokinesis and kept for 30h. Lastly, cells were fixed and stained for imaging. **(f)** Representative images of immunostaining on BN cells with MNi that are positive or negative for centromere staining (CREST) in non-treated (NT) condition. Phalloidin staining was employed to identify cytoplasms. White arrows indicate MNi. Cells were treated as indicated in panel (e) to block cytokinesis. Scale bar = 10 µm. **(g)** Representative images of immunostaining on BN cells with MNi that are positive or negative for centromere staining (CREST) in Mitomycin C (MitoC) treated condition. Phalloidin staining was employed to identify cytoplasms. White arrows indicate MNi.

Cells were treated prepared as indicated in panel (E) to block cytokinesis. Scale bar = 10 μm.

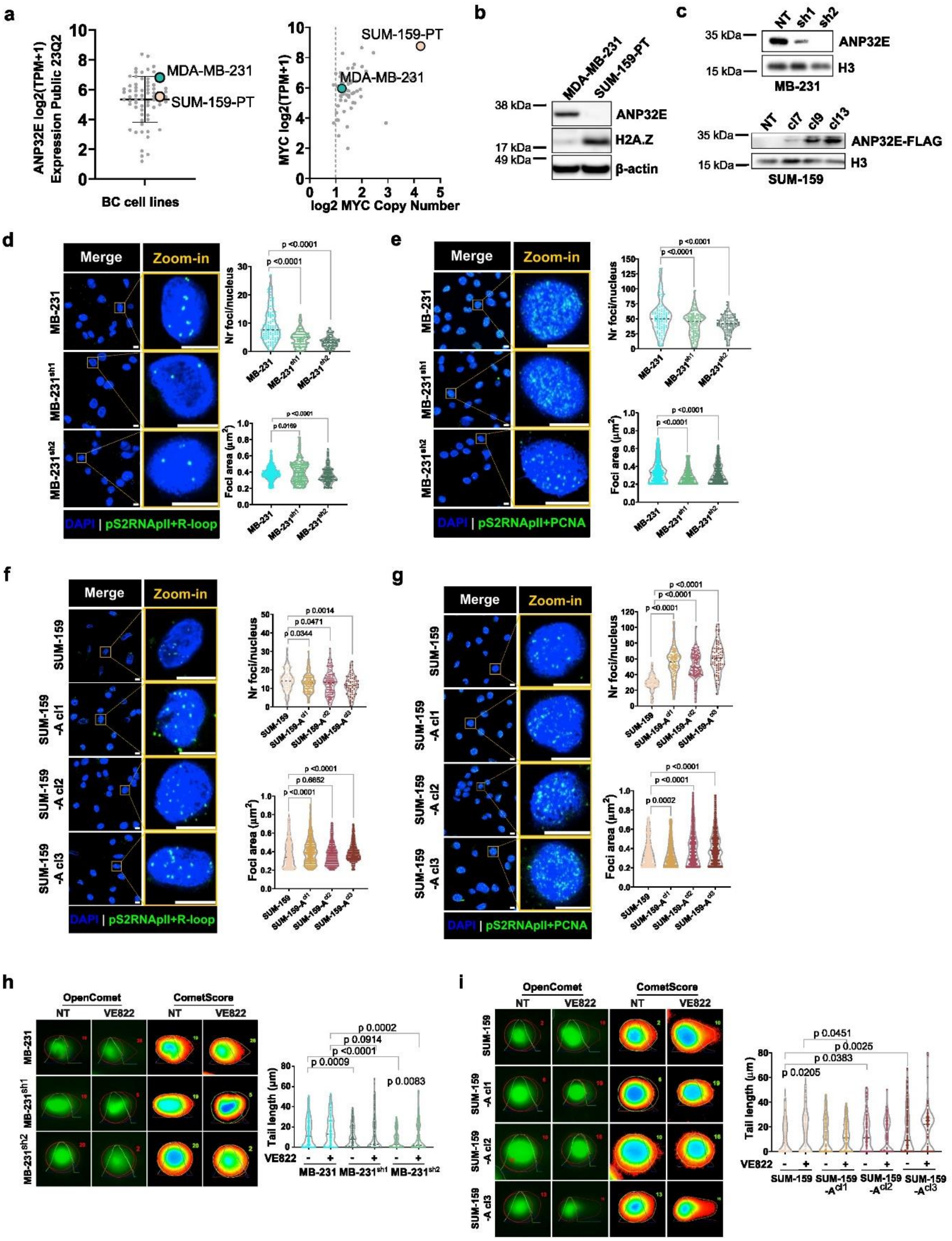


Lago_Extended Data Figure 3

**Extended Data Fig. 3. Validation of ANP32E-dependent stimulation of TRCs and DNA damage in alternative BC cell line models.**

**(a)** Left: Boxplot showing the mRNA expression level of ANP32E in BC cell lines retrieved from the DepMap portal in dataset 23Q2. MDA-MB-231 and SUM-159-PT are highlighted in the graph. Right: scatterplot representing MYC log2 mRNA expression in function of MYC log2 Copy number relative to ploidy +1 in BC cell lines retrieved from the DepMap portal in dataset 23Q4. MDA-MB-231 and SUM-159-PT are highlighted in the graph. **(b)** WB showing the protein level of ANP32E and H2A.Z in MDA-MB-231 and SUM-159-PT cell lines. B-actin is used as housekeeping for loading control. **(c)** Left: WB showing ANP32E silencing effect in MDA-MB-231 cells. H3 was used as housekeeping for loading control. Right: WB showing ANP32E overexpression in three different clones of SUM159-PT cells measured as FLAG-ANP32E. H3 was used as housekeeping for loading control. **(d)** Left: Representative images of PLA experiment between pS2RNApol II and R-loops in MDA-MB-231 cell lines with and without ANP32E silencing. Zoom-ins of single cells are reported. Scale bar = 10 μm. Right: quantification of foci number per nucleus and foci area. T-test p-values are indicated above comparisons for statistical significance. **(e)** Left: Representative images of PLA experiment between pS2RNApol II and PCNA in MDA-MB-231 cell lines with and without ANP32E silencing. Zoom-ins of single cells are reported. Scale bar = 10 μm. Right: quantification of foci number per nucleus and foci area. T-test p-values are indicated above comparisons for statistical significance. **(fF)** Left: Representative images of PLA experiment between pS2RNApol II and R-loops in SUM159-PT cell lines with and without ANP32E overexpression. Zoom-ins of single cells are reported. Scale bar = 10 μm. Right: quantification of foci number per nucleus and foci area. T-test p-values are indicated above comparisons for statistical significance. **(g)** Left: Representative images of PLA experiment between pS2RNApol II and PCNA in SUM159-

PT cell lines with and without ANP32E overexpression. Zoom-ins of single cells are reported. Scale bar = 10 µm. Right: quantification of foci number per nucleus and foci area. T-test p-values are indicated above comparisons for statistical significance. **(h)** Left: representative cells analyzed through Comet assay and comet profiles for MDA-MB-231 cells in the presence or absence of VE822 treatment. Right: Quantification of comets tail length. T-test p-values are indicated above comparisons for statistical significance. **(i)** Left: representative cells analyzed through Comet assay and comet profiles for SUM-159-PT cells in the presence or absence of VE822 treatment. Right: Quantification of comets tail length. T-test p-values are indicated above comparisons for statistical significance.

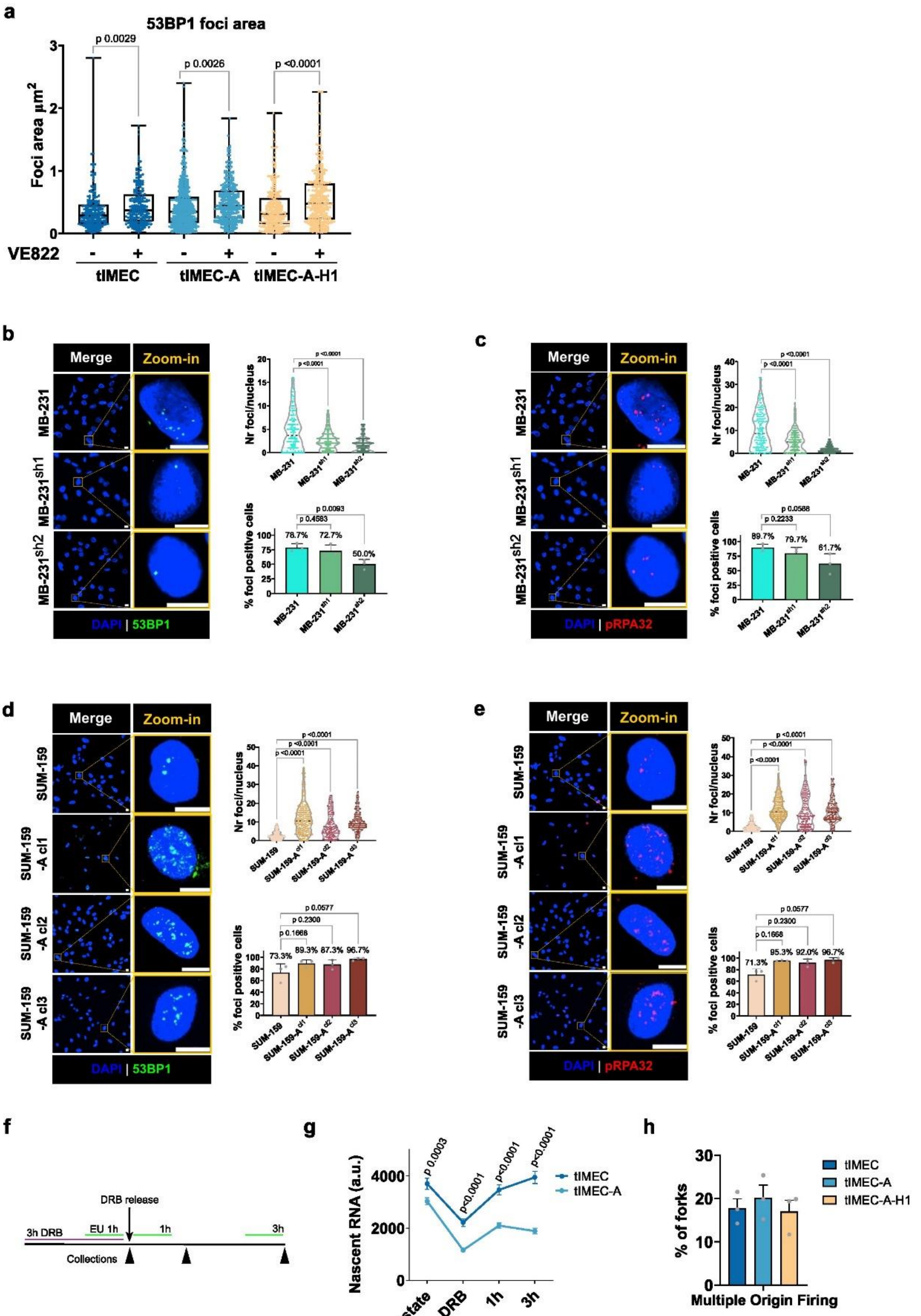


Lago_Extended Data Figure 4

**Extended Data Fig. 4. ATR-dependent DDR activation in ANP32E overexpressing BC cell lines.**

**(a)** Boxplot reporting the area of 53BP1 foci in untreated or VE822 0.25 μM 6h treatment condition. P-values for unpaired t-test are reported above the compared plots. **(b)** Left: representative images for 53BP1 immunofluorescence in MDA-MB-231 cells with and without ANP32E silencing. Scale bar = 10 μm. Right: quantification of foci number per nucleus and percentage of foci-positive cells. Unpaired t-test p-values are reported for statistical significance between comparisons as indicated. **(c)** Left: representative images for pRPA32 immunofluorescence in MDA-MB-231 cells with and without ANP32E silencing. Scale bar = 10 μm. Right: quantification of foci number per nucleus and percentage of foci-positive cells. Unpaired t-test p-values are reported for statistical significance between comparisons as indicated. **(d)** Left: representative images for 53BP1 immunofluorescence in SUM-159-PT cells with and without ANP32E overexpression. Scale bar = 10 μm. Right: quantification of foci number per nucleus and percentage of foci-positive cells. Unpaired t-test p-values are reported for statistical significance between comparisons as indicated. **(e)** Left: representative images for pRPA32 immunofluorescence in SUM-159-PT cells with and without ANP32E overexpression. Scale bar = 10 μm. Right: quantification of foci number per nucleus and percentage of foci-positive cells. Unpaired t-test p-values are reported for statistical significance between comparisons as indicated. **(f)** Scheme of DRB and EU treatments used to evaluate transcription dynamic at steady state, DRB block of elongating RNApol II enzyme and release. Lines length is proportional to the time of treatment. Purple indicate DRB treatment, green EU and black triangles indicate times of cells collection for EU staining and acquisition. **(g)** EU mean fluorescence intensity quantification. Mean and sd for every time point are reported. P-values for two-way ANOVA are reported above the compared points. **(h)** Percentage of DNA fibers that are classified as multiple origin firing according to the quantification of DNA fiber assay. Mean and sd of three biological replicates is reported. Statistic is not reported as not significant according to unpaired t-test p-values.

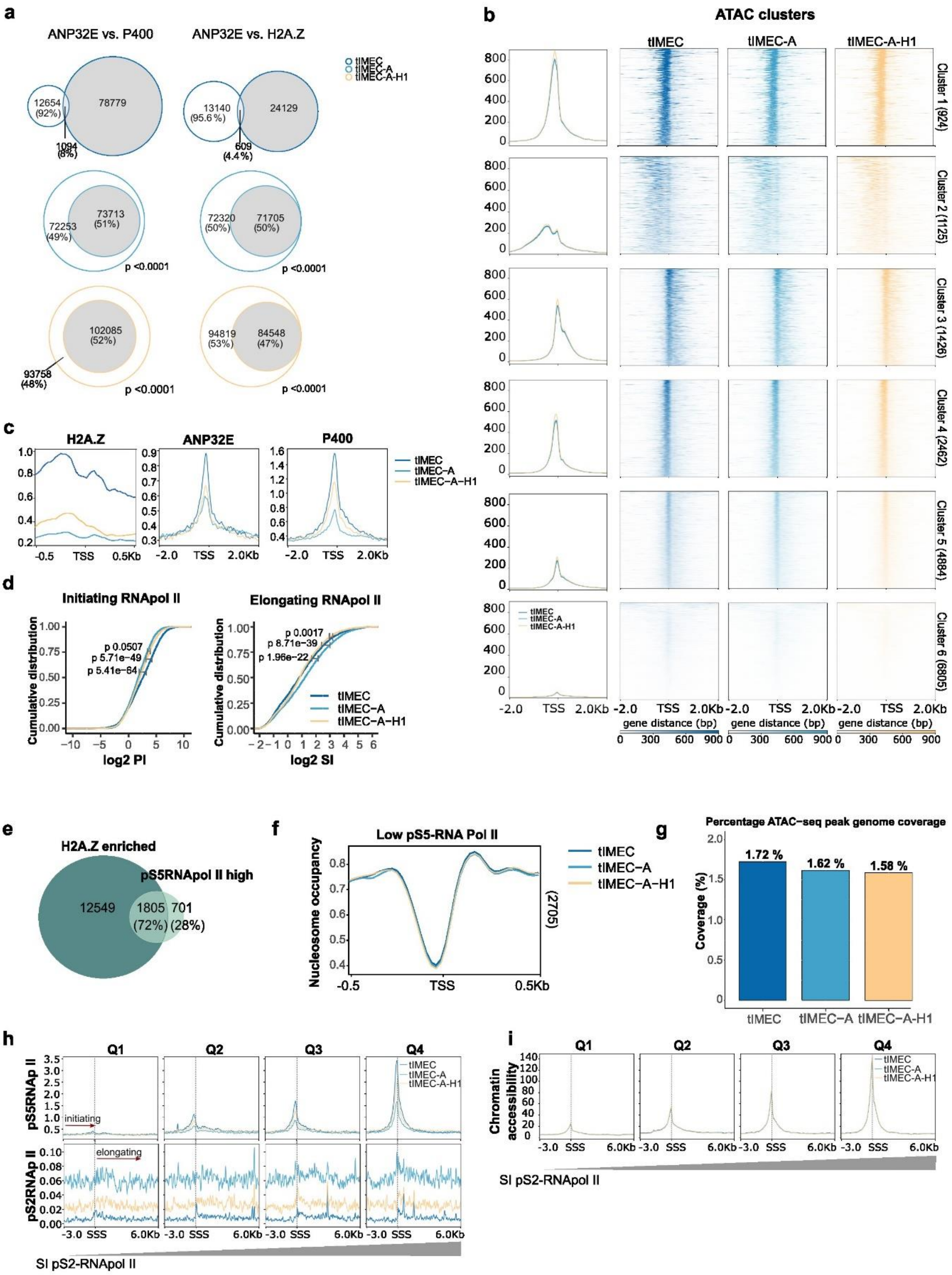


Lago_Extended Data Figure 5

**Extended Data Fig. 5. Genome-wide profiling of chromatin following ANP32E overexpression.**

**(a)** Venn diagrams reporting the percentage of peak overlap among ANP32E and p400 or H2A.Z. White-filled circles represent ANP32E, while grey-filled circles correspond to either p400 or H2A.Z, respectively. Percentages are calculated on the total number of peaks analyzed in each cell line. Peaks that are close to each other were merged into one. Hypergeometric test was used to calculate statistical significance of the overlap and is indicated for significant comparisons. **(b)** Heatmaps and profile plots of accessible promoters clustering into 6 clusters using k means algorithm for the three cell lines indicated in the legend. **(c)** Cumulative CUT&RUN signal plots of ANP32E, P400 and H2A.Z at pS5RNApol II-sorted ATAC-seq promoter clusters 1 to 5 (10,821 accessible sites). Heatmaps were plotted in genomic windows of ±2 Kbp for ANP32E and P400, and ±0.5 Kbp for. **(d)** Plot showing the PI of initiating pS5RNApol II (left) and the SI of elongating pS2RNApol II (right). Statistical significance p-values calculated by Wilcoxon test are reported for the indicated comparisons. **(e)** Venn diagram showing the overlap between peaks of the highest pS5RNApol II occupancy, and H2A.Z enriched region that have higher signal in tIMEC with respect to tIMEC-A. The percentage of pS5RNApol II peaks interesting or not those of H2A.Z is reported. **(f)** Nucleosome occupancy score cumulative plot at first (Low level) quartiles of pS5RNApol II in tIMEC accessible regions. The number of considered genomic regions is indicated on the right of each plot. **(g)** Barplot indicating the percentage of ATAC-seq peaks coverage in bp on the human reference genome for the indicated cell lines. **(h)** Cumulative plot of pS5- and pS2RNApol II on genomic regions of stalled pS2RNApol II grouped into quartiles according to the SI (n regions for every quartile = 2827). SSS indicates the Stalling Start Site. **(i)** Cumulative plot of ATAC-seq signal on genomic regions of stalled pS2RNApol II grouped into quartiles according to the SI (n regions for every quartile = 2827). SSS indicates the Stalling Start Site.

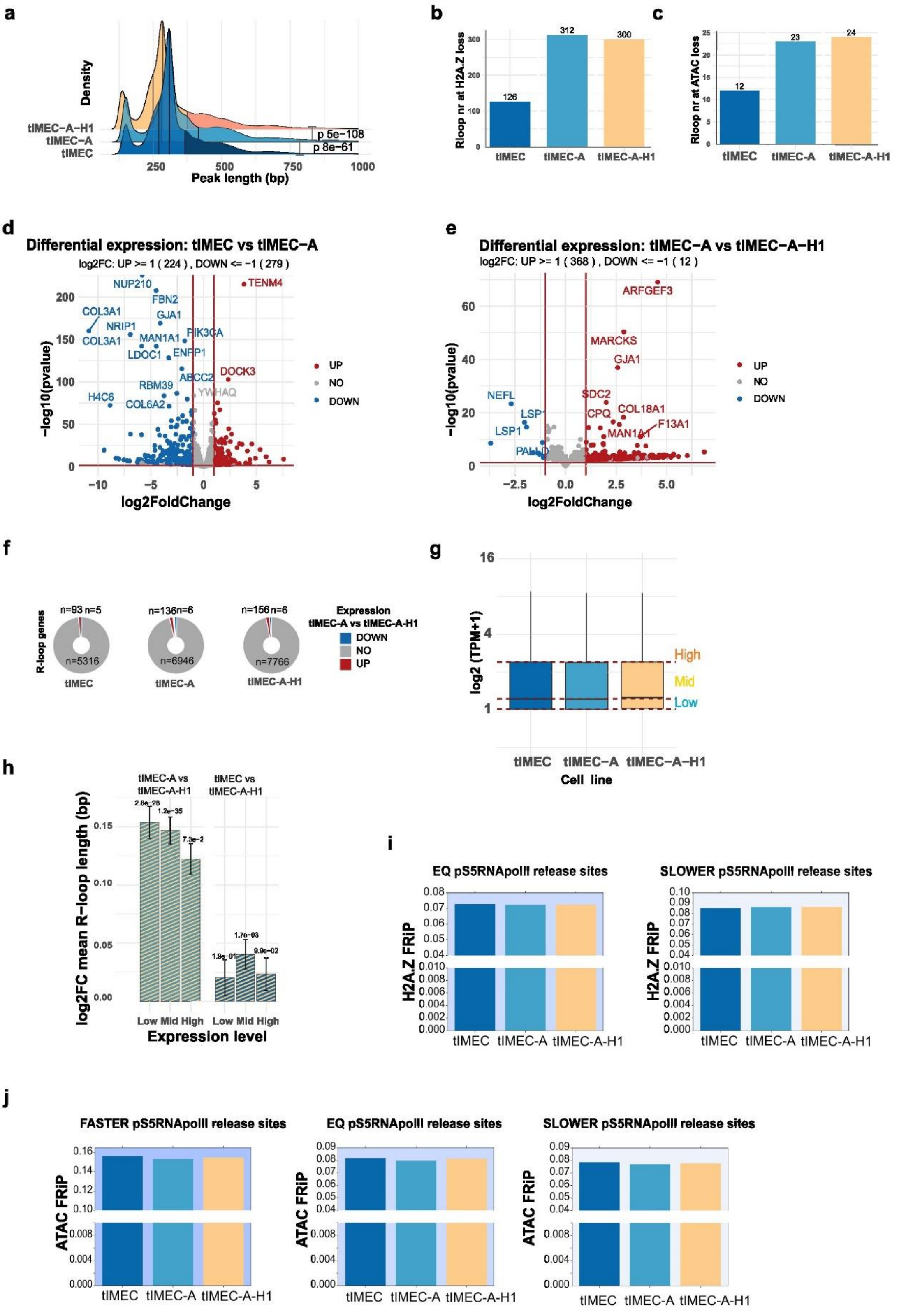


Lago_Extended Data Figure 6

**Extended Data Fig. 6. Evaluation of R-loop feature at genome-wide level upon ANP32E overexpression.**

**(a)** Density plot of R-loops in relation to their size in bp. The upper quartile (75°) is highlighted and t-test p-value reported on the graph. **(b)** R-loop peaks number at H2A.Z loss sites in tIMEC vs tIMEC-A. **(c)** R-loop peaks number at chromatin accessibility loss sites in tIMEC vs tIMEC-A. **(d)** Volcano plot of differential gene expression analysis for the comparison tIMEC vs tIMEC-A. **(e)** Volcano plot of differential gene expression analysis for the comparison tIMEC-A vs tIMEC-A-H1. **(f)** Pie charts reporting R-loop genes number that have differential gene expression (UP or DOWN) or are unvaried (NO) for tIMEC-A vs tIMEC-A-H1 comparison as calculated from RNA-seq data. **(g)** Boxplot showing the gene expression distribution for the cell lines tIMEC, tIMEC-A and tIMEC-A-H1. Genes with less than one transcript were considered not expressed. The remaining genes were divided in Low, Mid and High expression based on quartiles distribution in tIMEC as indicated on the graph. **(h)** Barplot reporting the mean and sd of the log2FC of differential R-loop length for the comparisons tIMEC-A vs tIMEC-A-H1 and tIMEC vs tIMEC-A-H1. R-loops were clustered in different groups based on the corresponding gene expression: Low, Mid, High. T-test p-values for statistical significance of the log2FC are reported above bars **(i)** Barplot for Fraction of Reads in Peak (FRiP) of H2A.Z in genomic regions where pS5RNApol II release is equivalent or slower in tIMEC-A vs tIMEC. **(j)** Barplot for Fraction of Reads in Peak (FRiP) of ATAC-seq data in genomic regions where pS5RNApol II release is faster, equivalent or slower in tIMEC-A vs tIMEC.

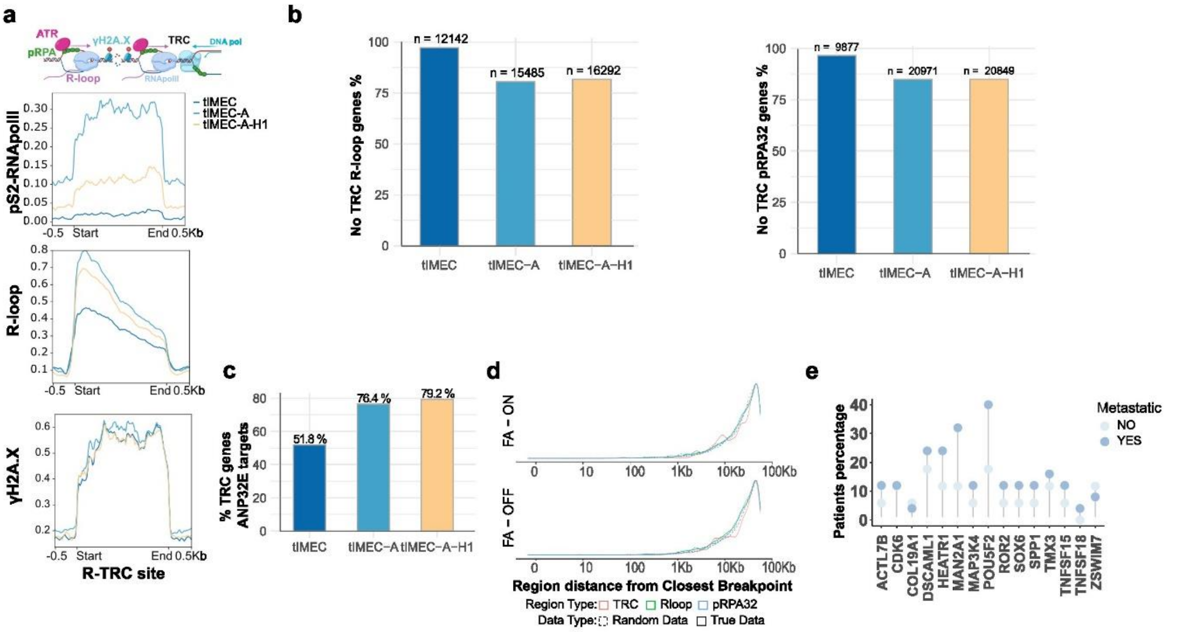


Lago_Extended Data Figure 7

**Extended Data Fig. 7. Evaluation of toxic TRCs features and potential association with tumor-related genome instability.**

**(a)** Cumulative plot of pS2RNApol II, R-loops and γH2A.X at toxic TRC regions (positive for pRPA32 and R-loops) scaled Start to End, with 0.5 Kb flanking. N regions = 7486. A scheme representing the proposed model of the TRCs is reported above the graph. **(b)** Bar plots reporting the percentage of genes in which non-toxic R-loops (left) or pRPA32-only were measured in each cell line. Raw number of genes is reported above each bar. **(c)** Bar plot reporting the percentage of TRC genes that are also ANP32E direct targets as measured by CUT&RUN data for the three indicated cell lines. **(d)** Density plot reporting the frequency of TRC regions (red) in relation to the distance from the closest Basal BC enriched CNV breakpoint in Basal BC patients stratified based on the activation of FA pathway, as derived from the RNA expression level of the genes belonging to the pathway. Regions characterized by non-toxic R-loops (R-loop only, green) or pRPA32 only (blue) are used as control regions as well as three sets of corresponding random regions (dashed lines). **(e)** Lollipop graph representing the percentage of patients with genomic alterations in genes that are actively expressed in our cellular model and are found in proximity of CFS breakpoints. Data retrieved from the Metastatic Breast cancer Project 2021.

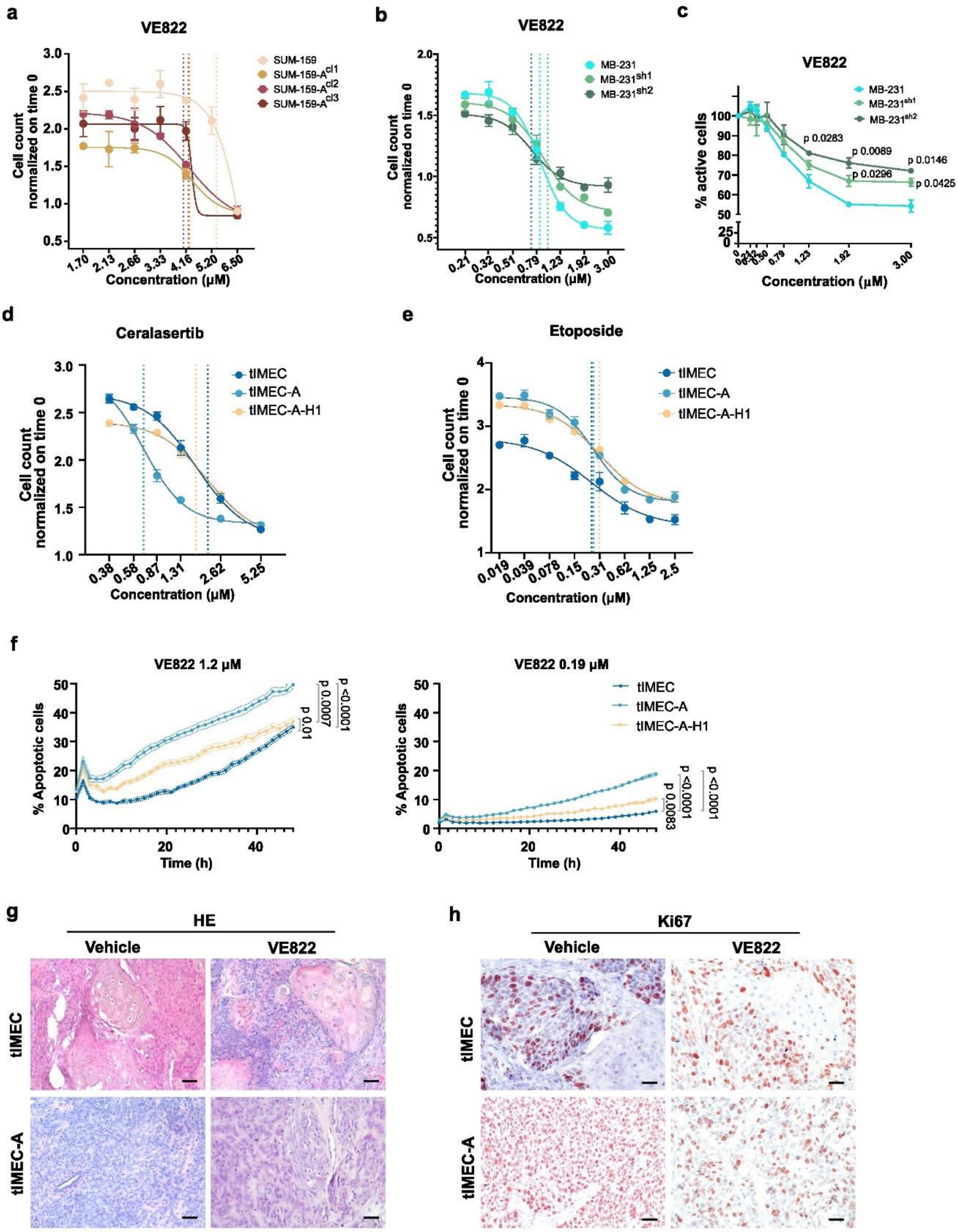


Lago_Extended Data Figure 8

**Extended Data Fig. 8. Evaluation of cytotoxicity and apoptosis induction in TNBC cell lines upon ATRi treatment. (a)** Representative drug response curves to 24h VE822 treatment of SUM-159-PT cells with and without ANP32E overexpression. Vertical dotted lines indicate the respective $EC_{50}$ value. **(b)** Representative drug response curves to 24h VE822 treatment of MDA-MB-231 cells with and without ANP32E silencing. Vertical dotted lines indicate the respective $EC_{50}$ value. **(c)** Viability curve measuring cellular metabolic activity upon 48h VE822 treatment of MDA-MB-231 with and without ANP32E silencing. **(d)** Representative drug response curves to 24h Ceralasertib treatment of tIMEC, tIMEC-A and tIMEC-A-H1 cell lines. Vertical dotted lines indicate the respective $EC_{50}$ value. **(e)** Representative drug response curves to 24h Etoposide treatment of tIMEC, tIMEC-A and tIMEC-A-H1 cell lines. Vertical dotted lines indicate the respective $EC_{50}$ value. **(f)** Percentage of apoptotic cells measured by caspase3/7 cleavage monitored for 48h upon VE822 treatment at a concentration higher (1.2 µM, left) or lower (0.19 µM, right) than the $EC_{50}$ values of tIMEC, tIMEC-A and tIMEC-A-H1 cells. **(g)** Hematoxylin and eosin staining of tissue sections obtained from tIMEC and tIMEC-ANP32E xenografts of mice treated with a vehicle or VE822. Scale bar 100 µm. **(h)** Ki67 IHC staining of tissue sections obtained from tIMEC and tIMEC-ANP32E xenografts of mice treated with a vehicle or VE822. Scale bar 100 µm.

#### TABLES:

**Supplementary Table 1**. Differential gene expression between MYC-CNA patients with Basal-BRCA (n=61) or Other-BRCA subtypes (n=92), retrieved from TCGA data.

**Supplementary Table 2.** 24h $EC_{50}$ Values for VE822, Ceralasertib and ETP treatments in the indicated cell lines.

**Supplementary Table 3.** List of Primers used for ATAC-seq library preparation.

**Supplementary Table 4.** List of antibodies used for CUT&RUN experiments.
